# Pathogen Host Shifts from a Niche Evolution Perspective

**DOI:** 10.64898/2026.08.31.748187

**Authors:** Peter Fransson, Henrik Sjödin

**Author notes:** Author for correspondence: Henrik Sjödin.

## Abstract

Emerging infectious diseases constitute a major health threat and are often associated with zoonotic spillover processes. Pathogen host shifts may involve evolutionary adaptation to new host species, which is of central concern in relation to future pandemics. Evolutionary host shifts require certain conditions to be met, including physiological and ecological factors. A pathogen strain infecting a new host species to which it is not well adapted will generally experience reduced success, which depends on the effective similarity between the new host species and the original reservoir host species from the pathogen’s perspective. The adaptation process involves intermediate strains, or phenotypes, that exist at the cost of reduced transmission and replication rates, and the evolutionary outcome depends on factors associated with host species physiology and population interactions. The interplay between host similarity and interaction, and pathogen adaptation trade-offs, is therefore important in steering pathogen evolution and possible evolutionary host-shift outcomes. Here we apply niche-evolution theory to host-pathogen systems to investigate the combined effect of host species similarity and intra- and interspecific interactions on evolutionary host shifts. We identify the conditions under which evolutionary host shifts occur and show, for a two-host-species system, that evolutionary outcomes fall into four distinct categories.

## 1. Introduction

Pathogen strains that emerge through transitions from its natural host to incipient host species constitutes a serious health threat for humans. It has been estimated that over 60% known human infectious diseases are zoonotic diseases and 75% of emerging diseases originate from non-human hosts [1,2]. Examples of such host shifts includes: HIV, where the pathogen jumped from non-human primates [3,4]; Influenza viruses like H1N1 (Spanish flue and swine flu) originating from birds and swine [5,6]; Coronaviruses like SARS-CoV (SARS) [7] and SARS-CoV-2 (COVID-19) [8]. Future projections predict an increase in pathogen jumps due to climate change driven host species distribution alteration [9].

While host shift events often result in severe healthcare burden, the emergence of pathogens that have not only transmitted through dead-end like spillover events, but are capable of standalone sustained transmission within the incipient host species, are relatively rare [10]. We will refer to this type of event as an evolutionary host shift. In evolutionary host shifts, the pathogen must overcome a number of barriers to be able to transmit from a reservoir host to and establish in an incipient host population. These include ecological and evolutionary barriers [10,11], which in turn can be organized across barriers at a higher level of detail (see e.g., [12]). From an ecological point, there needs to be opportunity for the pathogen to transmit from one reservoir host species to another incipient host species (Between-species barrier). Apart from a sufficiently high contact rate between the original and incipient host species populations, the contact rate between individuals of the incipient host species needs to be sufficiently high for the pathogen to spread in the novel host population (Within-species ecological barrier). This requires also that the incipient host species must also have physiological properties such that the pathogen can replicate within an individual of that species and further transmit to other individuals. From an evolutionary perspective, for new evolutionary strategies to survive (e.g., for host-shifts to realize), the pathogen’s original strategy in relation to the conditions for developing new ones, must be sufficiently beneficial in terms of fitness-tradeoffs following Darwinian principles. Thus, both host-species ecology (such as host species distribution overlap and interspecific interaction) and similarity between host species (for example the phylogeny of host species) are important factors for the occurrence of zoonotic spillover and determining the success of host shifts [13,14]. However, the complexity and impact of host species similarity and interspecific interaction on pathogen evolution and evolutionary host shifts is an area which requires further research [15].

Considering the importance of understanding how different factors impact pathogen evolution and host shifts, a number of modeling studies have focused on zoonotic diseases and pathogen emergence. [10] employed a within-host, individual-based (virion-level), model with a genetic algorithm to simulate the transmission dynamics between a reservoir species host and a novel species. The model used an explicit representation of the pathogen’s genome, in the form of a binary array (array of zeros and ones). The study revealed that the between-host barrier significantly hindered the establishment of adapted strains in the novel species, with increasing fitness slope and fitness differences negatively impacting virus emergence. [16] employed an extended SIR (Susceptible-Infectious-Recovered) model to simulate pathogen transmission from a wild reservoir host to humans, involving an intermediate domestic species. The infection network included interactions where infected wild hosts could transmit to both susceptible wild hosts and domestic hosts. Additionally, infected domestic hosts could infect susceptible domestic hosts and humans, while infected humans could only transmit to susceptible humans. The study considered two pathogen strains, a wild-type and a mutant-type, with the mutant-type arising from a mutation in a wild-type infected domestic host. The findings indicated that a pathogen with a basic reproduction number (R_0_) less than one could be maintained in the human population if allowing for mutation within an intermediate host. [17] introduced an evolutionary game theory-based SIS model involving two species, a reservoir host and an incidental host, with a focus on pathogen strains. The study assumed a fixed number of strains in the reservoir population, where only one strain can jump to an incidental host. Strain competition was represented by a weighted directed graph, depicting the potential for one strain to superinfect another. The researchers analyzed a three-strain system using a rock-paper-scissors framework, revealing cyclic frequency patterns under specific conditions.

The concept of a pathogen having a focal host species, the infection success within a novel host species depending on the similarity between them [14], and the trade-off caused by host shifting is a type of niche evolution in the context of consumer-resource models in theoretical ecology [18–21]. These models assume that different types of consumers compete for resources of varying types. In an evolutionary context, the consumer strategy, e.g., the focal resource type of a consumer (the niche), is dynamically changing through eco-evolutionary feedback. Thus, such a framework appears promising to apply to evolutionary host shift scenarios; however, none of the modelling studies mention above have explicitly incorporated such a framework in a host population-level in their work.

Here we introduce an epi-evolutionary framework in a consumer-resource niche evolution context to investigate how the combined effect of host species similarity and intra- and interspecific contact rates impacts pathogen evolution of pathogens and evolutionary host shifts. Specifically, we introduce a multi-host-species-and-multi-strain stochastic SIS where transmission parameters are determined by 1) the host species (resource type) and 2) the trait value of the pathogen (consumer strategy). To study the evolution of our model, we apply a framework of deterministic approximations of individual-based stochastic processes referred to as adaptive dynamics [22–24]. While our framing and model study are described with a virus in mind, many of the described mechanisms fit generally also to other types of pathogens, yet important variations exist. For this reason, while not claiming it fits to all types of pathogen, nor that it is streamlined to any specific, we will for practical purposes use the general term “pathogen” when we refer to the infecting agent in this study.

## 2. Methods and theory

### (a) The resource space and the *τ*-function

In theoretical ecology, consumer-resource models are a class of models in which a collection of consumers compete for resources of varying types. Typically, the consumers have different levels of adaptation to the resource types, e.g., they have a varying ability of acquiring the resource types. In an evolutionary setting, the consumer strategy, e.g., focal resource type, is assumed to be a heritable [19]. Within our modelling framework, we treat pathogen strains as consumers and host species, or, more accurately, susceptible cells within individuals of the host species, are considered resources. To model the pathogen adaptiveness to a host species, the pathogen-host combability, we assume that host species are represented by points in a predefined resource space ℛ(Figure 1). ℛcould for instance represent a specific set of host phenotypes and other predictors linked to the host-pathogen interaction, such as host specific covariate, e.g., host body size, host geographical range, and climate factors at host habitat, and host-host covariates (dyadic covariate), e.g., phylogenetic distance between host species [13,25–27]. Further, we assume that such a distance between points is a measure of similarity between resources (from the perspective of the pathogen), i.e., host species that are more similar (i.e., more likely to share pathogens) have point representations closer to each other. We will refer to this distance as the host-species similarity. Strains differ with respect to how closely adapted they are to the respective hosts. The degree of adaptation to any given host species is quantified by the variable *τ*, the probability of infection conditional on an interaction between a susceptible and infected individual that could lead to an infection. Thus, we have a *τ*-value for every strain-host pair; more specifically, for every strain, we have a *τ*-value for every point in ℛ. Furthermore, we assume that there exists one point in ℛ where a strain experiences the largest *τ*-value among all points in ℛ, and, in a sense, the point to which it is best adapted. This point is called the niche position [19]. Pathogen strains differ in terms of this point (i.e., each strain is uniquely defined by its niche position) and we assume that the niche position is a heritable trait. Additionally, we assume that *τ* declines monotonically around the niche position. The *τ*-value will therefore typically decrease with distance between the host and the focal host of the strain in question (i.e., the niche position). Let *d*(*x, y*) denote the distance between *x, y* ∈ ℛ. Then, assuming that the two different host-species are defined at *x* and *y*, respectively, the distance *d*(*x, y*) is equivalent to the host-species similarity. If we denote the position of a given novel species by *x* ∈ ℛ, and a focal species that a specific strain is perfectly adapted to by *z* ∈ ℛ, we can define *τ* as,

**Figure 1.**
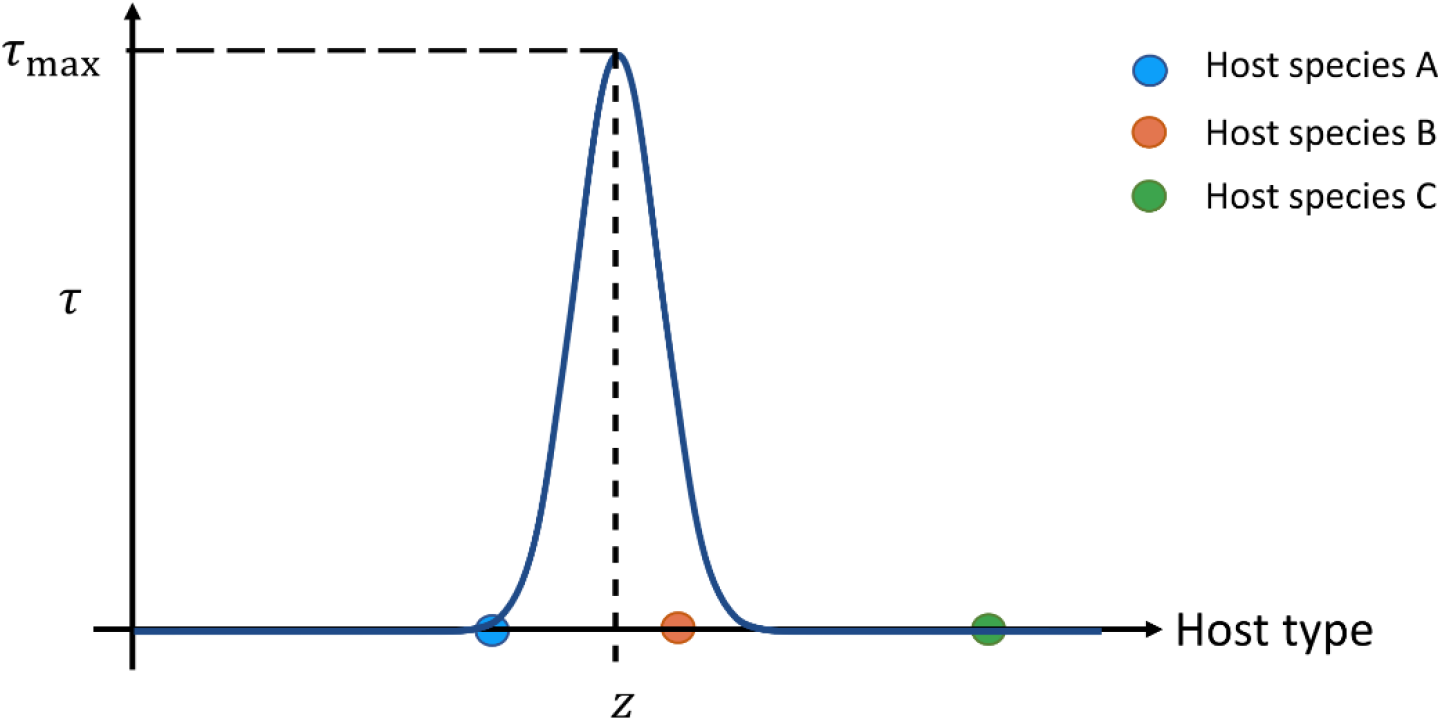
Host-pathogen combability. We assume that host species are represented as points in a resource space (the set of host types). Here, the resource space is represented by the x-axis. A pathogen strain is defined by the niche position, z (a position in resource space). z represents the specialization/natural host of the strain. The combability between a strain and a host species is quantified by the probability of infection (τ), conditional on a pathogen-host contact. A specific strain has its highest combability, τ_max_, with a species located at z. Compatibility between the strain and host species decreases monotonically with the distance between the point representation of the host species and z (host-species similarity). The blue curve represents the compatibility between a particular pathogen strain and all possible host types (the x-axis), and the three colored points are representations of the host species a, b, and c.

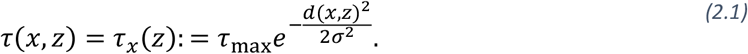

Here *τ*_max_ is the maximum attainable *τ*-value and *σ* is the niche width, a measure of how fast *τ*(*x, z*) declines with *d*(*x, z*). A graphical representation of *τ*(*x, z*) is provided in Figure 1.

### (b) Stochastic epi-evolutionary model

For our analysis, we apply a multi-host-species-and-multi-strain stochastic SIS (Susceptible-Infected-Susceptible) model. The basic, classical, SIS model describes a system where a pathogen is transmitted from infected individuals (*I*) to susceptible individuals (*S*), and previously infected individuals do not acquire long-term immunity after recovery, thus becoming susceptible again. The dynamics of the basic SIS is described by the following system of ordinary differential equations (ODE),

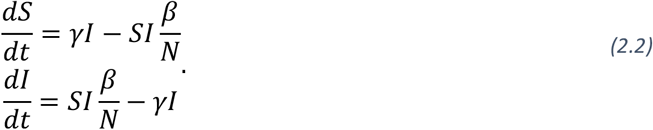

Here, *N* is the total population size, *γ* is the recovery rate, *β* = *τc* is the transmission rate, *τ* is the conditional probability of infection, and *c* is the contact rate. The system can reach an endemic state if the basic reproduction number, *R*_0_ = *β*/*γ*, is greater than one, while the system has only the disease-free state as an equilibrium when *R*_0_ is less than one. The basic SIS model can be derived as a deterministic approximation of the stochastic SIS model [28]. In the context of our stochastic model, the basic SIS model would represent the deterministic approximation of a system with one host species and one evolutionary inert pathogen strain. We will show here how we set up a more complex multi-species and multi-strain stochastic SIS that allows for exploration of strain evolutionary processes in the context of between host-species adaptation, i.e., evolutionary host shifts.

We will now describe the stochastic individual-based epi-evolutionary SIS model (Figure 2); state variables and parameters are listed in Table 1. The model assumes a fixed number of interacting host individuals, where each individual belongs to one of *n*_*s*_ ∈ ℕ host species. Further, the individuals can be in one of two states, susceptible (*S*_*i*_) or infected (*I*_*i,z*_). Here, *i* ∈ {1, ⋯, *n*_*s*_} denotes the species, and the infected state can be further subdivided based on the strain that an individual is infected by, which is denoted by *z* ∈ ℛ (the pathogen strain trait value). In contrast to the basic SIS model, *S*_*i*_ and *I*_*i,z*_ are integers rather than continuous real numbers. The state of an individual (susceptible, infectious, the strain that infects the host) changes over time through three types of events: 1) Infection, *S*_*i*_ + *I*_*j,z*_ → *I*_*i,z*_ + *I*_*j,z*_. 2) Recovery, *I*_*i,z*_ → *S*_*i*_. 3) Mutation, 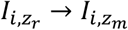. Here *z*_*r*_ denotes the current resident strain trait value which the individual is infected with and *z*_*m*_ is a mutant strain trait value.

**Table 1.**
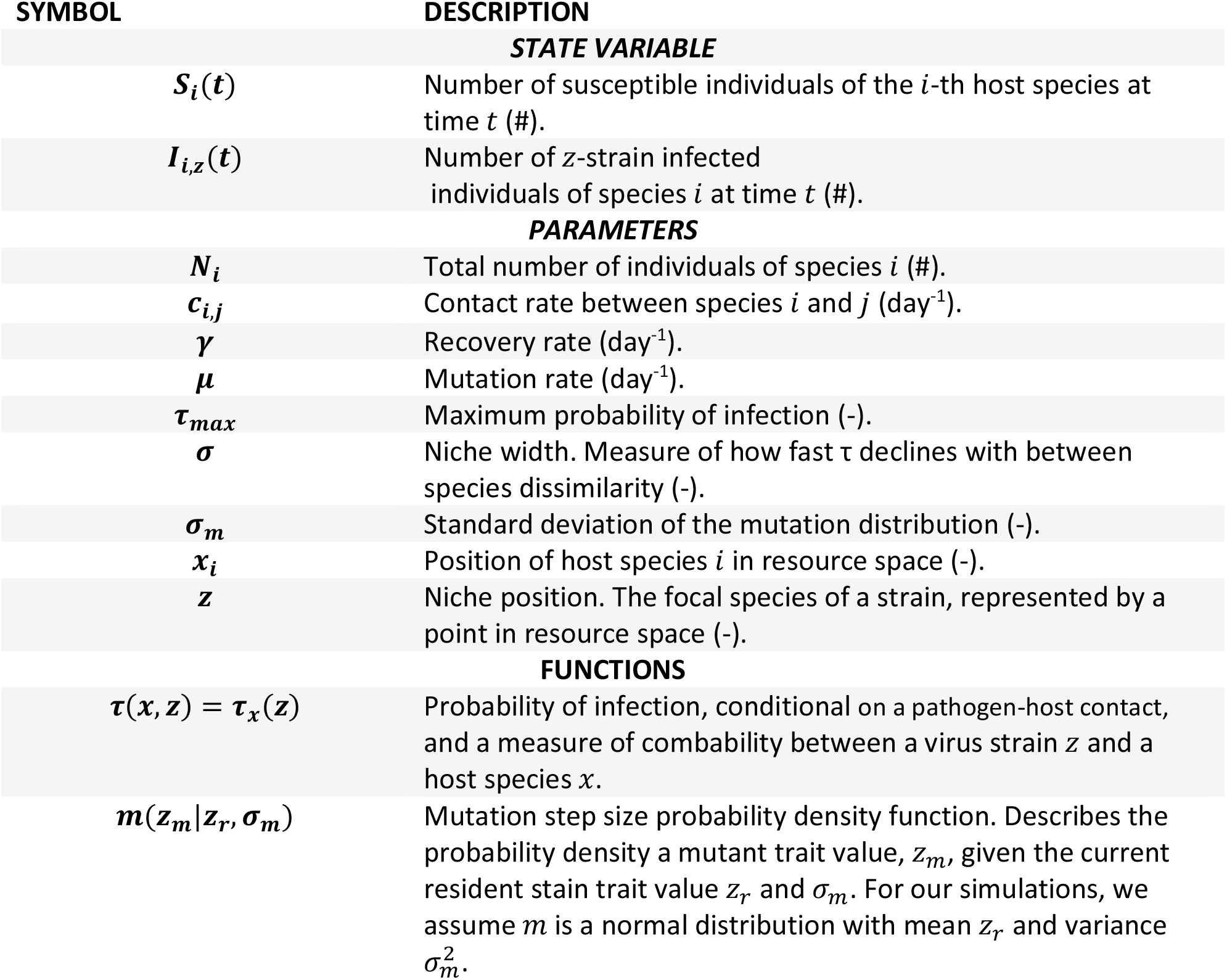
State variables and parameters of the stochastic individual-based model. *Units are given in parentheses*.

**Figure 2.**
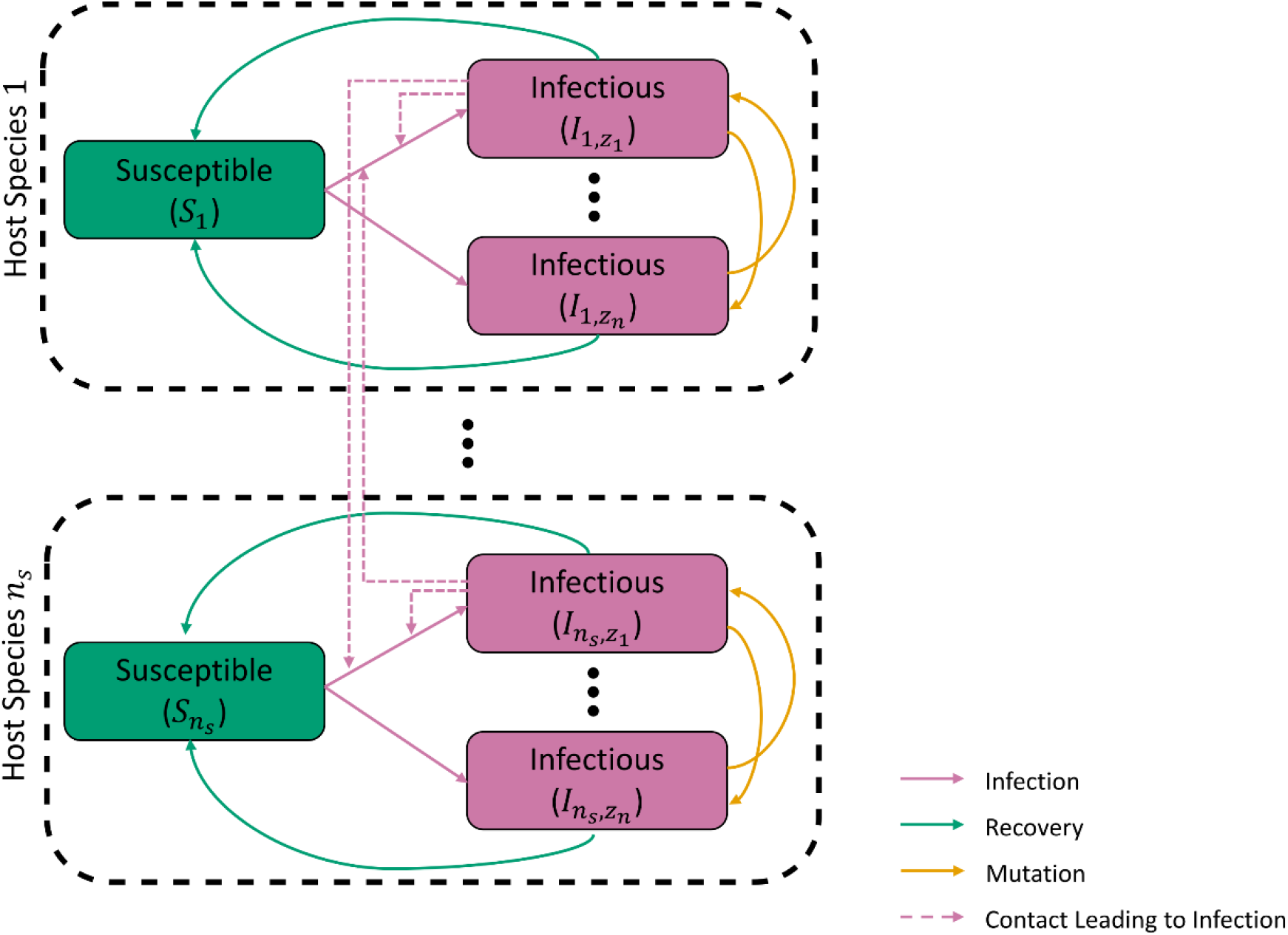
The modelled epi-evolutionary SIS (Susceptible - Infectious - Susceptible) system comprises of n_s_ interacting host species. Each species population, i, is divided into susceptible (S_i_) and infectious individuals 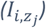, and the total population of i is indicated by the inside of a dashed rectangle. The infectious population is further subdivided, depending on the pathogen strain causing the infection. The specific strain is denoted by the continuous index z in I_i,z_. Sizes of the subpopulations 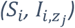 changes dynamically due to: 1) Infections resulting from contacts between susceptible and an infectious individual (inter- and intraspecific contact, marked with dashed lines in the figure). Note, for the sake of brevity we have restricted to only displaying interactions that lead to an z_1_ infection in species 1 and n_s_ in the figure. 2) Recovery of an infectious individual. 3) Mutations within infectious individuals, giving rise to new pathogen strains. The probability (density) of emergence of a new strain is conditional on the present resident strain. Note, the number of strains circulating in the system n, changes over time. The total host populations are assumed to be constant.

There are eight fundamental assumptions underpinning our model: i) Individuals and events (infection, recovery, mutation) are independent. ii) Each individual is infected by at most one strain at a time (no co-infection and superinfection assumption) [29]. iii) Infected individuals give rise to a mutant strain with rate *μ*. iv) mutant strain trait values (*z*_*m*_) are drawn from a mutation step size distribution (*m*(· | ·)) centered around the current resident strain trait value (*z*_*r*_) and standard deviation *σ*_*m*_, i.e., *z*_*m*_∼*m*(*z*_*m*_|*z*_*r*_, *σ*_*m*_). v) Infected individuals recover at a rate *γ*. vi) The transmission rate between a susceptible individual of species *i* and an individual of species *j*, infected by a strain with trait value *z*, is equal to *τ*_*i*_(*z*)*c*_*i,j*_, where *c*_*i,j*_ denotes the contact rate between species *i* and *j*. vii) Interspecific contact rates are symmetric (*c*_*i,j*_ = *c*_*j,i*_). viii) Population dynamics of host species populations are not considered (no births or deaths among the host populations) and *N*_*i*_ denotes the total number of individuals in the *i*-th host-species population. With these eight assumptions, the system can be modeled as a continuous-time Markov jump process.

The system has two types of state variables, the number of susceptible individuals of the *i*-th host species at time *t* (*S*_*i*_(*t*)), and the number of *i*-th host species individuals infected with strain *z* ∈ ℛ at time *t* (*I*_*i,z*_(*t*)). Sample paths can be generated from this process by applying the classic Gillespie Stochastic Simulation Algorithm (SSA), also referred to as the Direct Method [30]. In short, the time between events, Δ*t*, is an exponentially distributed random variable, i.e., Δ*t*∼Exp (∑ _*e*_ *α*_*e*_(***x***_*t*_)).

Here, ***x***_*t*_ is the current state of the system which is defined by the number of susceptible individuals and the strain distribution (the collection of *I*_*i,z*_) for each species, at time *t*, and *α*_*e*_ is the propensity of event *e*. The equations to calculate *α*_*e*_ for any event *e* is listed in Table S 1 (Supplementary material S1). The probability of a specific event *e, p*(*e*), is proportional to *α*_*e*_, i.e., *p*(*e*) = *α*_*e*_(***x***_*t*_)/ ∑ _*e*_ *α*_*e*_(***x***_*t*_). To speed up computation, we apply the hybrid algorithm proposed by Zhu et al. ([31]). Zhu et al. applied their hybrid algorithm to a population genetics model of cancer development. The efficiency of the hybrid method relies on the combined employment of the exact SSA for critical events, such as mutation events, and the approximate tau-leaping (not to be confused with the variable *τ*(*x, z*) in our study) algorithm [32] for non-critical events, see Supplementary material S1 for more information.

### (c) Evolutionary invasion analysis

We apply methods from the adaptive dynamic framework [22,23,33] –also referred to as evolutionary invasion analysis– to study the evolutionary outcomes of pathogen strains in our stochastic individual-based SIS model. In this section we will only give a brief overview of the most relevant parts of the framework, for a more comprehensive introduction we refer to the paper by [34], but see also Supplementary material S2. The framework can be used to study the trait evolution. A core assumption is that the change in trait values (evolutionary dynamics) of the strains and the transmission dynamics act on different timescales. Specifically, evolutionary dynamics are assumed to be sufficiently slow in relation to “demographic” dynamics. This warrants the assumption of separation of timescales in mathematical models, i.e., we can assume the population under evolution reaches a steady state before a new mutant strain with trait value *m*, appears in the system. Important to note in the context of our study on pathogen evolution, while the time-scale separation assumption per se implies long-term evolutionary dynamics, it must be seen from the perspective of the evolving unit, and we must interpret “long-term” in relation to the fast transmission and mutation rate of pathogens, meaning that pathogen-evolution should be expected to be relatively fast in relation to our perception of epidemic or endemic progressions.

The steady state is dependent on the current trait value, *r*, of the resident strain in the pathogen population. The success of rare strains with trait value *m* is dependent on its expected initial per capita population growth rate in a steady state resident population. This is captured by the invasion fitness *S*_*r*_(*m*). More specifically, *S*_*r*_(*m*) is the expected per capita growth rate of a strain with trait value *m* when it is initially rare within the steady state population with trait value *r*, thus *S*_*r*_(*m*) is a function of both *m* and *r*. The initially rare mutant population with trait value *m* has a chance to successfully invade if *S*_*r*_(*m*) > 0 or will go extinct if *S*_*r*_(*m*) < 0. Per definition *S*_*r*_(*r*) = 0.

To calculate *S*_*r*_(*r*), we derive the system of ordinary differential equations (ODE) (Eq. S 1) which represents the deterministic approximation of the expected dynamics of our stochastic model, see Supplementary material S2. The ODE is derived using mean field approximation [35] for the one-strain-multi-host version of our stochastic SIS model, i.e., the case when only one strain is present in the system and mutation events are absent. The resulting ODE system has the form,

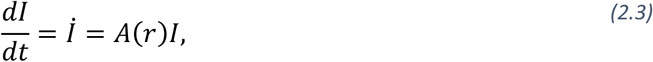

where, 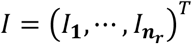 is the vector of infected individuals for all *n*_*r*_ number of host species and *A*(*r*) is the system matrix Eq. S 2. Note, *A*(*r*) is dependent on the resident strain trait value *r* ∈ ℛ. From this ODE, we can determine the endemic state (steady state), i.e., *İ* = **0**. Additionally, we also use the ODE to derive the expected dynamics for the rare mutant *m* ∈ ℛ (Supplementary material S2). The ODE that describes the expected initial growth of *m* has the same form as (2.3), but the system matrix is now a function of both *m* and *r, A*(*m*|*r*) (Eq. S 3). Based on stability theory for ODEs, we define the invasion fitness as 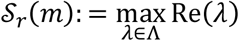, where Λ is the spectrum (set of eigenvalues) of the system matrix *A*(*m*|*r*), see Supplementary material S2. The selection gradient can be calculated by taking the derivative of *A*(*m*|*r*) with respect to *m* and multiplying the resulting gradient matrix by the left and right eigenvector associated with the eigenvalue *λ*, see Supplementary material S2 for more information. Note, as an alternative to *S*_*r*_(*m*), we can also use the reproductive number of *m* at the endemic state of *r*, ℜ(*m*|*r*). The quantity ℜ(*m*|*r*) − 1 has the same sign as the invasion fitness *S*_*r*_(*m*) [36], and ℜ(*m*|*r*) is also calculated from *A*(*m*|*r*) via the next-generation matrix procedure [37].

We can derive a lot of insight by studying *S*_*r*_(*m*) and its derivatives. The direction of evolution can be determined by the derivative,

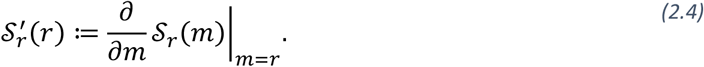

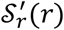 is called the selection gradient and when 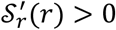 then the resident population can be invaded by a mutant strain with slightly higher trait values; the opposite is true when 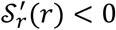. Traits, *r*^∗^, where the selection gradient vanishes, i.e., 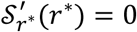, are of particular interest since these signify a potential endpoint of evolution. These traits are known as evolutionary singular strategies, and one can classify them by investigation, using higher order derivates. We call *r*^∗^ an evolutionarily stable strategy (ESS) if 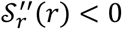, i.e., if *r*^∗^ is a local fitness maximum. However, if 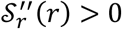 then *r*^∗^ is a local fitness minimum at which disruptive selection –evolutionary branching– can occur [34]. If small perturbations in *r*^∗^ lead to strategies with selection gradients guiding the trait evolution back to *r*^∗^, then *r*^∗^ is called a convergence stable strategy, mathematically this is equivalent to,

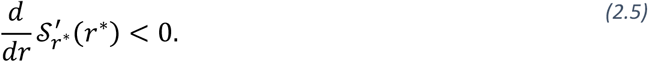

We can graphically visualize the evolutionary dynamics by so-called pairwise invasibility plots (PIPs). These plots are dichromatic heatmaps where the color is determined by the sign of *S*_*r*_(*m*) for a given *r* and *m* pair, the x- and y-coordinate, respectively. From these plots we can also find any singular strategy, where the boundary of the positive *S*_*r*_(*m*) region intersects the diagonal (the *r* = *m* line), and classify them [34].

### (d) Case study: The dual host-species system

We explore and analyze the long-term evolutionary outcome for the dual host-species system, host species *a* and *b*, using adaptive dynamics. For this case we assume that ℛ ∈ ℝ, the host-species similarity between the two species is calculated as *d*(*x, y*) = |*x* − *y*|, and *x*_*a*_ and *x*_*b*_ denotes the position of species *a* and *b* in resource-space respectively. Further, without loss of generality, we assume *x*_*a*_ < *x*_*b*_. We perform a parameter exploration and record every unique class of evolution outcome. The parameter ranges are a provided in Table 2 and we apply Sobol sequences –quasi-random numbers– to generate samples of parameter combinations to efficiently and evenly fill out the parameter space, similar to [38]; For our parameter exploration we used 10^6^ samples. We classify the evolutionary outcome in terms of: 1) the number of singular strategies, 2) their individual classification (ESS and/or convergence stable), and 3) if the system *R*_0_ is above 1 for all resident strain strains *x* ∈ [ *x*_*a*_, *x*_*b*_].

**Table 2.**
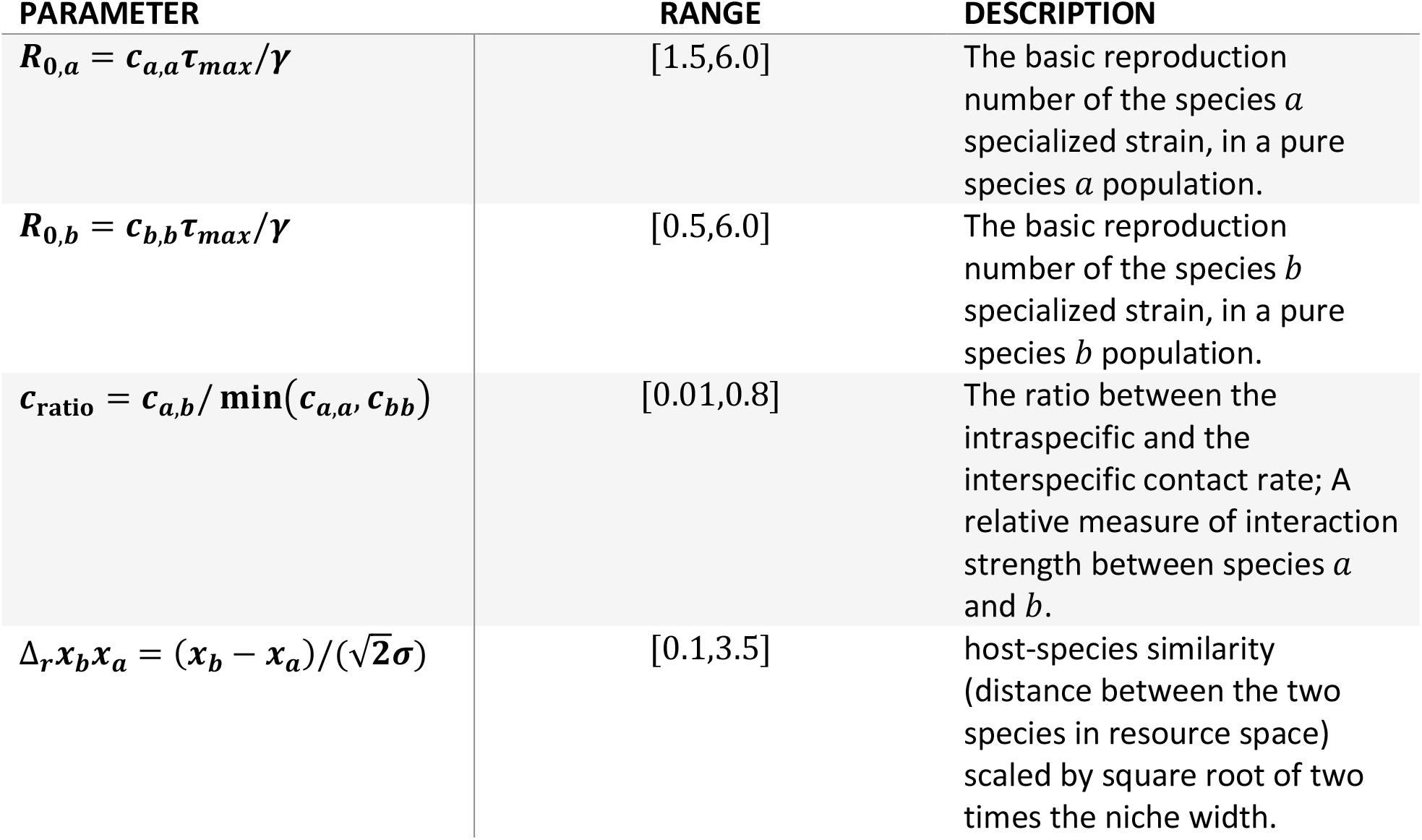
Parameter ranges used in the parameter space exploration.

| PARAMETER | RANGE | DESCRIPTION |
| --- | --- | --- |
| $R_{0,a} = c_{a,a}\tau_{max}/\gamma$ | [1.5,6.0] | The basic reproduction number of the species $a$ specialized strain, in a pure species $a$ population. |
| $R_{0,b} = c_{b,b}\tau_{max}/\gamma$ | [0.5,6.0] | The basic reproduction number of the species $b$ specialized strain, in a pure species $b$ population. |
| $c_{ratio} = c_{a,b}/\min(c_{a,a}, c_{b,b})$ | [0.01,0.8] | The ratio between the intraspecific and the interspecific contact rate; A relative measure of interaction strength between species $a$ and $b$. |
| $\Delta_r x_b x_a = (x_b - x_a)/(\sqrt{2}\sigma)$ | [0.1,3.5] | host-species similarity (distance between the two species in resource space) scaled by square root of two times the niche width. |

Additionally, we investigate which parameter combinations describe a potential evolutionary host shift scenario (EHSS) and how many of these scenarios lead to an evolutionary host shift (EHS), under small mutation assumption (*σ*_*m*_ approaches zero). To this end, we define EHSS and EHS, together with some auxiliary terms, in the context of this study. We call a strain trait value, *z*, adapted and maladapted to the *i*-th host species if the basic reproduction number of *z* in a pure *i* population is >1 (*R*_0_(*x, z*) = *c*_*i,i*_*τ*(*x*_*i*_, *z*)/*γ* > 1, where *x*_*i*_ is the position of *i* in ℛ) and <1, respectively. We define EHS, with respect to the *i*-th host species, as the event when a *i*-adapted strain, *z*, emerges, and can persist long-term, in the system, when initially no species *i*-adapted strains were present in the system. The definition implies that: 1) evolutionary processes are needed for the EHS to occur (for *z* to emerge) and 2) that *z* does not require interspecific interactions to continuously transmit in the pure *i* population. From this definition, it is clear that not all parameter combinations describe a system where an EHS has the potential to occur. For the dual host-species system we have two crucial conditions for EHS, with respect to *b*: 1) *z* = *x*_*b*_ is adapted to *b*, and 2) the strain *z* = *x*_*a*_ is maladapted to *b*. This ensures that we have *b* adapted and maladapted strains in *x* ∈ [ *x*_*a*_, *x*_*b*_]. We define an evolutionary host shift scenario (EHSS), for the dual host-species system with respect to species *b*, as a parameter combination which fulfills both conditions. This allows us to study the evolution of an initially monomorphic strain population *z* = *x*_*a*_ (wild-type strain), when its natural host (*a*; reservoir host) comes into contact with an incipient host species *b*, and if the trait evolution leads to an EHS (with respect to *b*).

For the special dual host-species case we derived explicit expressions for 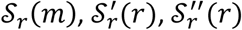, and the endemic state of the deterministic one-strain SIS model, which are provided in Supplementary material S3. We use a symmetric difference quotient approximation to calculate the necessary derivative (2.5), to determine convergence stability. To find all singular strategies, we applied the *find_zeros* function from the *Roots*.*jl* package [39].

## 3. Results

### (a) Evolutionary outcome is determined by four parameters

From analysis of the dual host-species system, we found that we can reparametrize the model and reduce the number of parameters need to draw PIPs from ten to four, see Supplementary material S3. Specifically, for the dual host-species system, with host species *a* and *b*, the parameters needed to draw a PIP are: 1) *R*_0,*a*_ = *c*_*a,a*_*τ*_*max*_/*γ*, the basic reproduction number of the species *a* specialized strain, in a pure host species *a* population. 2) *R*_0,*b*_ = *c*_*b,b*_*τ*_*max*_/*γ*, the basic reproduction number of the species *b* specialized strain, in a pure host species *b* population. 3) *c*_ratio_ = *c*_*a,b*_/ min(*c*_*a,a*_*c*_*b,b*_), a relative measure of interspecific interaction. 4) 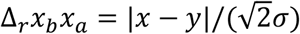, the scaled host-species similarity between host species *a* and *b*.

### (b) The four major outcome-types of long-term evolution

From the parameter exploration of the dual host-species system, we found that the evolutionary outcome of the vast majority of the sample points, 99.7% (997328 out of 10^6^), will fall into one out of four evolutionary-outcome types (Figure 3). These types vary in terms of the number and quality of evolutionary singular strategies: (1) Generalist outcome (375145 out of 10^6^ ≈ 37.5%). In this case there exists a single evolutionary singular strategy, in between the two host species, which is convergence stable and ESS, i.e., the strain adapts to both species (Figure 3 a). (2) Evolutionary branching outcome (122168 out of 10^6^ ≈ 12.2%). Here there exists one evolutionary singular strategy in ℛ, which is convergence stable strategy, but not evolutionary stable (Figure 3 b). This leads to an evolutionary branching event where the initially monomorphic strain population undergoes disruptive evolution and splits into a dimorphic population. A trait evolution plot (TEP) [22] reveals that the dimorphic population has an evolutionary singular strategy in the region of coexisting strains (Figure S4) close to the optimal strategy of both species. Thus, the long-term evolutionary outcome results in a system with two coexisting strain trait values, one adapted to *a* and one to *b*. (3) Bistable outcome (201341 out of 10^6^ ≈ 20,1%). For this type, we get three evolutionary singular strategies in ℛ (Figure 3 c). Two convergence stable and ESS, located close to the two host species. Additionally, we have an intermediate strategy which is both convergently and evolutionary unstable. The two ESS are not global maximums; thus, the evolutionary outcome is stochastic. Specifically, we either observe the population of circulating strain trait values getting stuck in one of the ESS (depending on our initial state), or we observe a branching event (Figure S 3). The probability of a branching event is highly depending on the assumptions of the mutation process, e.g., the mutation rate, variance of the mutant distribution, and the distribution itself, etc. Under the small mutation assumption, the evolution ends in the local optimum. (4) Isolated bistable (298674 out of 10^6^ ≈ 29,9 %). Similar to the bistable outcome, we have two convergence and evolutionary stable strategies, one for each species. However, instead of having a non-convergent stable and non-ESS singular strategy between the two ESSs, there exists a zone of intermediate trait values for which *R*_0_ < 1, for the dual host-species system. Thus, the strategies in these zones only have the disease-free state as an equilibrium state. Beside the four aforementioned types, we also found intermediate type (2672 out of 10^6^ ≈ 0.27%). These only appear rarely as transitional type, between the main types (Figure 4), and the evolutionary outcome appears to mainly fall into the four categories outlined above.

**Figure 3.**
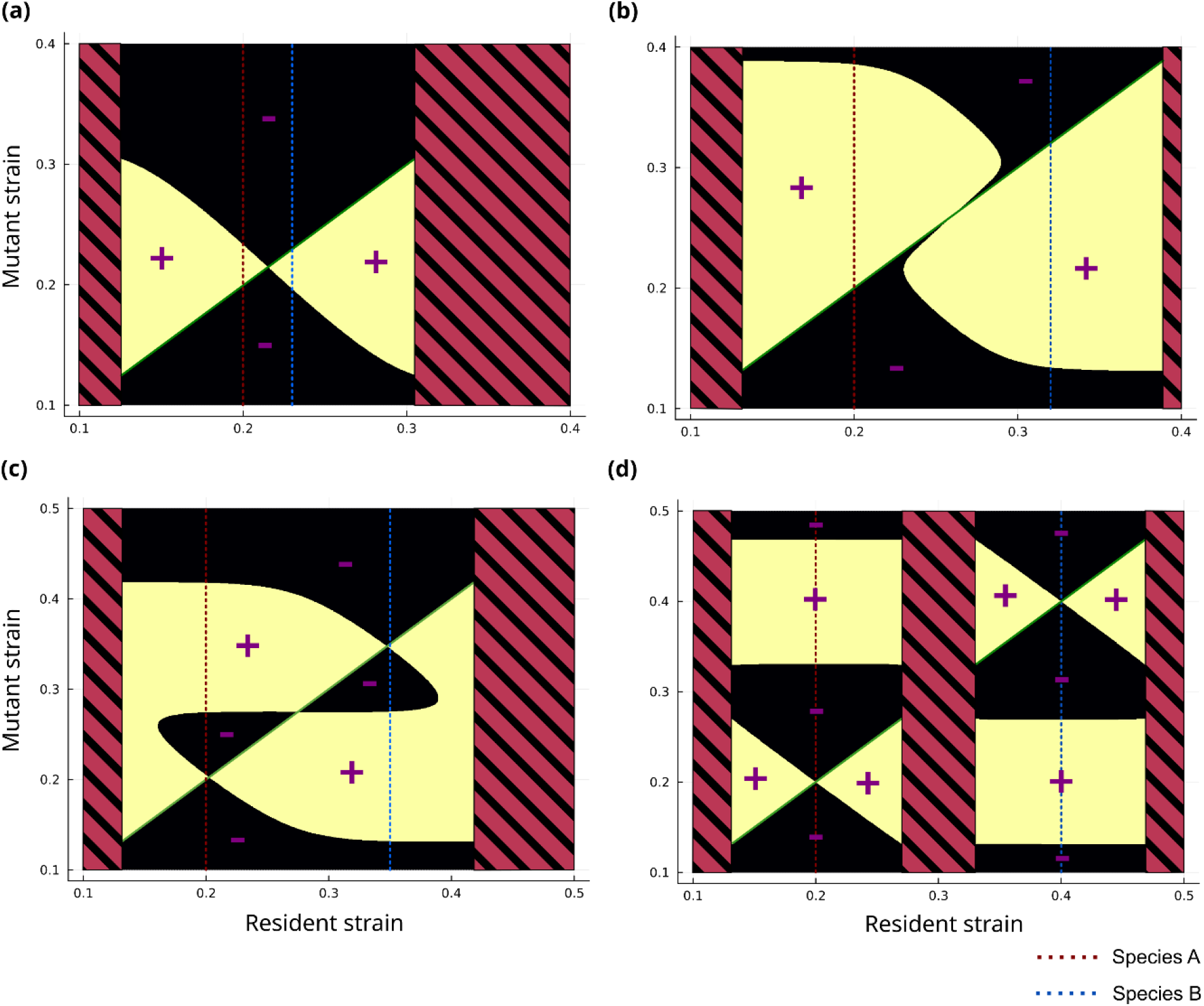
Pairwise invasibility plots for the dual host-species system. Areas marked with plus-signs and minus-signs indicate when a mutant strain trait value has a positive invasion fitness (S_r_(m) > 0) and negative invasion fitness (S_r_(m) < 0), respectively. The striped area indicates resident strains for which the disease-free state is the only stable equilibrium, i.e., the basic reproduction number (R_0_) of the system is less than one. The host-species similarity between host species A and B is given by their distance in the recourse space (ℛ) (graphically depicted as the distance between the red and blue dashed vertical lines). We have four main evolutionary outcomes. Generalist (a): One evolutionary singular strategy which is convergence stable and ESS. Evolutionary branching (b). Bistable (c): three evolutionary singular strategies, two of these are convergence stable and ESS and the third is neither convergence stable or ESS. Isolated bistable (d): similar to the bistable outcome but there exists a region of intermediate resident strains where R_0_ < 1. The figures were generated using the following parameter values: R_0,a_ = R_0,b_ = 2.55, c_ratio_ = 0.71, and Δ_r_x_a_x_b_ = 0.42 (a), = 1.75 (b), = 2.12 (c), = 2.83 (d).

**Figure 4.**
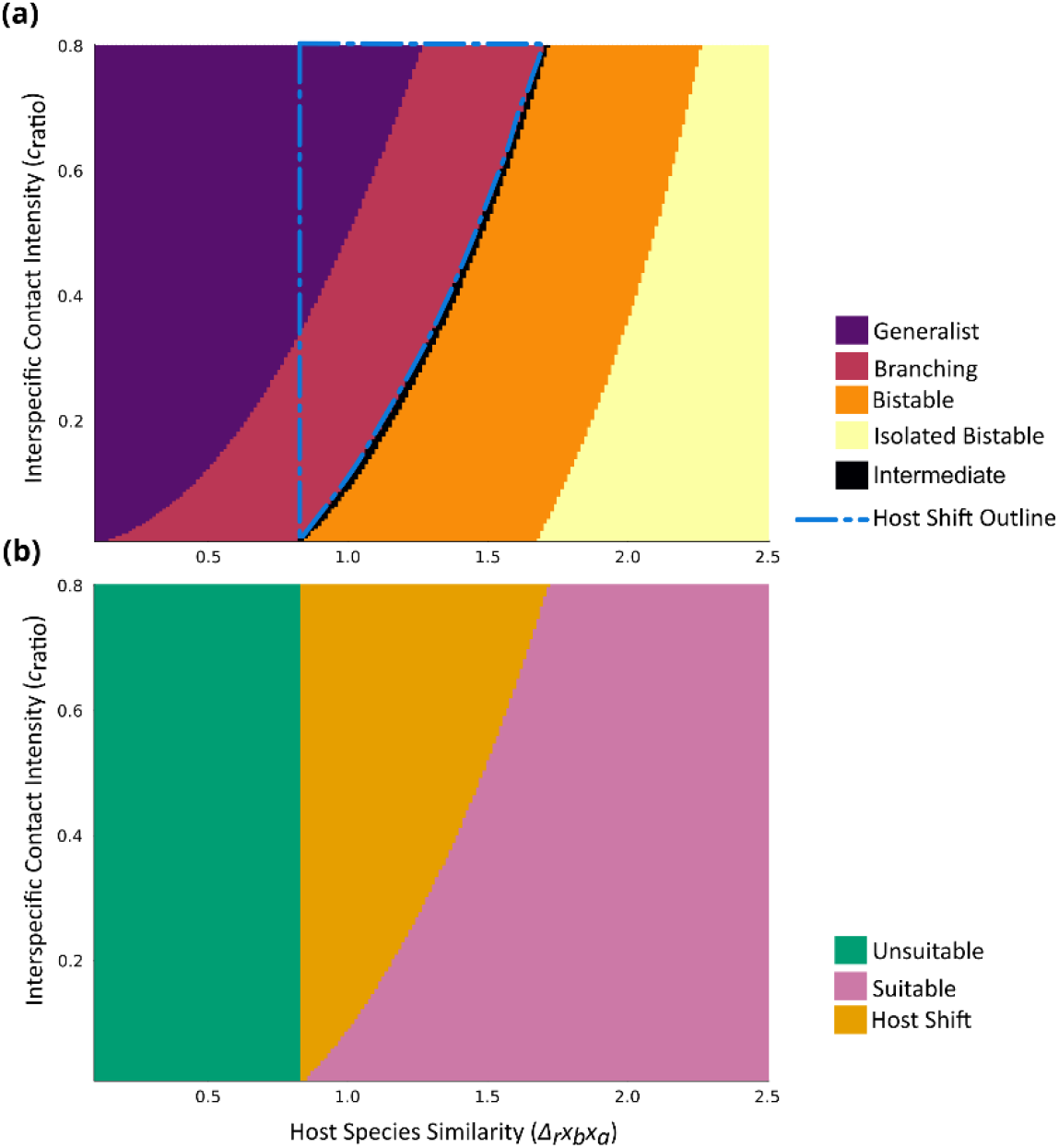
The interspecific contact intensity (c_ratio_) impacts the evolutionary outcome and evolutionary host shift (EHS) events. The color of the top heatmap (a) indicates the evolutionary outcome (Generalist, (evolutionary) Branching, Bistable, Isolated Bistable, and Intermediate outcomes) for different combinations of host species similarity (Δ_r_x_b_x_a_) and c_ratio_. The color of the bottom heatmap (b) indicates if a Δ_r_x_b_x_a_ and c_ratio_ combination describes a system which is: not an evolutionary host shift scenario (EHSS) (Unsuitable), i.e., a parameter combination not fulfilling necessary conditions for a EHS to occur. EHSS, but EHS did not occur (Suitable). EHHS and EHS occurred (Host Shift). Note the dashed curve in (a) indicates the corresponding outline of the Host Shift region in (b). Both heatmaps where generated using R_0,a_ = R_0,b_ = 2.0.

As a robustness check, we conducted simulations, using the stochastic SIS model for the generalist (Figure S 1), evolutionary branching (Figure S 2), and bistable (Figure S 3) outcome in Supplementary material S1. The stochastic simulations verified that the deterministic findings are generally are robust.

### (c) Interspecific contact intensity changes the evolutionary outcome and evolutionary host shift events

Interspecific contact intensity (*c*_ratio_) has an impact on the evolutionary outcome. From our simulations we find that for a given between host-species similarity (Δ_r_*x*_b_*x*_a_), *c*_ratio_ will shift evolutionary outcome (Figure 4 a). For example, in Figure 4 a, when Δ_r_x_b_x_a_ = 1.0, lower *c*_ratio_ will result in a bistable outcome, while intermediate and high *c*_ratio_ results in evolutionary branching and generalist outcome respectively. Likewise, *c*_ratio_ impact the range of Δ_r_x_b_x_a_ for which EHS occurs (Figure 4 b). Specifically, we found that higher *c*_ratio_ resulted in a wider Δ_r_x_b_x_a_ range for which EHS occurs, i.e., higher interspecific contact intensity allows for host shifts between more dissimilar host species. In Figure 4 we assumed *R*_0,*aa*_ = *R*_0,*bb*_, we provide additional figures in the supplementary material section (Figure S 5 in Supplementary material S5) where we investigate the impact of the *R*_0,*bb*_/*R*_0,*aa*_ ratio. We found that an increase in *R*_0,*bb*_/*R*_0,*aa*_ increases the minimum Δ_r_*x*_b_*x*_a_ for which EHS occurs and an increase in Δ_r_*x*_b_*x*_a_ and *c*_ratio_ combinations resulting in EHS. Additionally, we also found that the *R*_0,*bb*_/*R*_0,*aa*_ ratio will impact the evolutionary outcome. A decrease *R*_0,*bb*_/*R*_0,*aa*_ results in a reduction in Δ_r_*x*_b_*x*_a_ and *c*_ratio_ combinations resulting in evolutionary branching and below a critical ratio value we find that no combinations result in a branching event. An increase in *R*_0,*bb*_/*R*_0,*aa*_ will also increase the number of Δ_r_*x*_b_*x*_a_ and *c*_ratio_ combinations resulting a bistable outcome.

### (d) Only a small proportion of potential evolutionary host shift opportunities are realized

Recall that for potential evolutionary host shifts to realize, it is required that: 1) There exist at least one strain in ℛ (the pool of all potential strain trait values) which is *b*-adapted and 2) The wild-type strain (*z* = *x*_*a*_) is *b*-maladapted. To check if the two requirements are fulfilled and consequently if a given parameter combination is an EHSS, with respect to *b*, we have to test: 1) 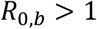 and 2) 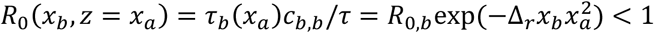. Note the second condition implies that *b* and *a* cannot be too similar, i.e., 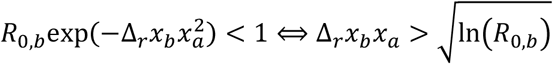. We found 65.8% (658185 out of 10^6^) of the samples where EHSS. Additionally, we checked how many of the suitable parameter combinations result in an EHS, with respect to *b*, given sufficient time. This can be determined by the PIPs. More specifically, we get an EHS if the EHSS parameter combination has either an evolutionary branching outcome or the evolutionary singularity closest to *x*_*a*_ is convergence stable and ESS, where the ESS strain trait value, *z*_ESS_, fulfils the condition *R*_0_(*x*_*b*_, *z*_ESS_) > 1. We found that 14.8% of the EHSS parameter combinations (97102 out of 658185) resulted in an EHS. Most, 64.3% (62413 out of 97102), of the EHS occur as a consequence of an evolutionary branching and the remaining 35.7% (34689 out of 97102) from a generalist outcome.

## 4. Discussion and conclusion

### (a) Model scope and limitations

Our model and framework integrate concepts from ecological and evolutionary consumer-resource theory (e.g., [19], allowing us to incorporate the impact of host species similarity and pathogen host specificity on transmission parameters in a host population-level epidemiological compartmental model. This is of significant interest since host similarity (e.g. phylogenetic relativeness), particularly for RNA viruses, and between species transmission are known important predicter and bottleneck of viral sharing among mammal host spices [10,14,40]. In our model, natural selection is purely driven by the trade-off in *τ*(*x, z*)-values for a pathogen strain and the considered host species. This allows us to study the evolution of pathogen populations under different scenarios (different parameter combinations). We studied the single and combined effect of host species similarity and interspecific interaction, and which model parameter combinations results in an evolutionary host shift.

Our results showed that only a low number of potential evolutionary host shift scenarios are actually realized. This means that, among the systems (parameter combinations) in which an evolutionary host shift is possible, only a few experiences an evolutionary host shift due to the direction of natural selection. However, the estimated approximately 15% realized evolutionary host shift scenarios should be viewed as a lower bound estimate. Less restrictive assumption on mutation would result in a higher proportion evolutionary host shift realization. For example, our result showed that no bistable outcome resulted in an EHS (Figure 4 and Figure S 5), under small mutation assumption, but in our stochastic simulations we observed EHS for a bistable system when allowing for larger *σ*_*m*_ (Figure S 3). In fact, if there is a non-zero probability (density) of emergence for all pathogen strains, then evolution would eventually lead to a branching event for the bistable and isolated bistable case, conditioned on that the total pathogen population doesn’t go extinct. Thus, the assumptions regarding the mutation step-size probability (density) function (*m*(· | ·)) highly influence strain emergence. We assumed *m*(· | ·) follows a normal distribution centered around the resident strain, thus allowing for the emergence of incipient host adapted strains in the incipient host. This is sometimes referred to as “tailor-made” emergence [10]. Other, more restrictive assumptions like “off-the-shelf” emergence only, i.e., novel host adapted strains only emerges in the reservoir host, would likely influence our results. However other studies have shown that both “tailor-made” and “off-the-shelf” emergence scenarios showed similar behavior with respect to the relation between infection probability in a novel host and the between host fitness difference [10]. Additionally, the parameter ranges impact the percentage of realized EHS.

The focus of our study was the combined effect of host interaction and pathogen-host compatibility on pathogen evolution and host shifts. As such we do not account for processes affecting host population dynamics like virulence, and natural birth and death process. These could be readily introduced in future studied by, for example, imbedding our framing work to a population dynamics model like the model initially developed by Doobs [41–43]. However, introducing this process would result in a similar system matrix (Eq. S 2); the big difference would possibly manifest in the equilibrium points. Additionally, we have assumed that the inheritable trait in our system is continuous. This should be regarded as an approximation of general pathogen–host systems. In reality, changes in traits may take discrete forms, such as changes in receptor structure, particularly in specific pathogen–host systems (see, e.g., [44]).

### (b) The impact of interspecific interaction and host species similarity on pathogen evolution and host shifts

Our results showed that, for a fixed between host similarity (Δ_*r*_*x*_*b*_*x*_*a*_), that the increase in interspecific interaction (*c*_ratio_) can lead to an evolutionary host shift when *c*_ratio_ is above a critical threshold level (Figure 4). This is in line with previous findings showing that actions and processes altering habits of wildlife, including wildlife exploitation, land use changes, other anthropogenic activities, and climate change are important predictors of pathogen spillover risk [9,13,45]. This is due to the increase in human-animal interaction and creation of novel species-spices interactions. While we restricted to the case of static contact rates, these processes are realistically dynamic, and would in the context of our model imply time dependent contact rates and contact network structure. This would be of interest to incorporate and study in the future, yet require a mathematically more advanced and complex approach.

When fixating interspecific host interaction (i.e., keeping *c*_ratio_ constant), the overall impact of host-species similarity is that evolutionary host shift occurs in a range from a lower to an upper host-species similarity limit (Figure 4). If Δ_*r*_*x*_*b*_*x*_*a*_ is sufficiently small, the initial pathogen strains in circulation in the reservoir host (host *a*) are already adapted to the novel host (host *b*) and doesn’t require any adaptation (“off-the-shelf” emergence). However, if the two host species are sufficiently dissimilar (Δ_*r*_*x*_*b*_*x*_*a*_ is sufficiently large), then the *τ*-value is too small for a reservoir-host-adapted strain infecting the novel host, and evolutionary intermediate populations, that would be required for a branching event, is not viable and is not realized. As discussed previously, the upper host similarity limit depends on the assumption of the mutation distribution. However, even for wider mutation distributions (larger *σ*_*m*_) the probability of EHS (occurring during a given time period) decreases with Δ_*r*_*x*_*b*_*x*_*a*_. This is caused by the increase in the required host-species distance between mutant and resistant strain (see ESS in Figure 3 C and D). While these are expected properties of such an evolutionary system, we have here to show a quantification of them for evolutionary host shifts. [13] showed similar result, where they found that increased phylogenetic dissimilarity between host species generally results in lower viral sharing probability (probability of two species sharing at least one pathogen) across different degrees of habitat overlap.

The combined of effect of the degree of interspecific interaction (i.e., *c*_ratio_) and host-species similarity (i.e., Δ_*r*_*x*_*b*_*x*_*a*_) manifest in: 1) Increased interspecific interaction increases the range of host-species similarity, for which we observe EHS. Thus, allowing for EHS for more dissimilar host species pair. 2) That the threshold level of *c*_ratio_ increases with increased Δ_*r*_*x*_*b*_*x*_*a*_. This means, that a higher level of interspecific interaction is required for an EHS for more dissimilar host species pair. This shows the importance of interspecific interactions, such as wildlife exploitation and hunting, for the emergence of pathogens and have to be considered when designing intervention strategies.

### (c) The impact of the dimension of the resource space

In our analysis, we only considered a one-dimensional resource space (ℛ). The dimension represents a covariate or predictor of the host-pathogen combability, e.g., the phylogenetic distance in the two-host species case. The extension of ℛ to higher dimension can be done in a relatively straightforward way by allowing ℛ to extend from ℝ to ℝ^*n*^ and using the Euclidian distance as the proxy for host-species similarity, see Supplementary material S6. The interpretation of a higher dimensional resource space is that the host-pathogen combability is explained by more than one covariate, e.g., host phenotypes and traits, and predictors linked to the host-pathogen interaction [26,27]. It should be noted, when extending ℛ to ℝ^*n*^ and applying the Euclidean distance as a measure of host-species similarity, it involves a number of implications. In particular, the units of the underlying covariates are assumed to be comparable, i.e., one-unit difference in one covariate is equal to a one-unit change in the others, and that the underlying covariates assumed to be uncorrelated. If the covariates are correlated, one could construct new, independent composite covariates, for example, by applying principal components analysis, see Supplementary material S6 for an extended discussion.

The high-dimensional ℛ introduces new pathways and directions for strain evolution, compared to the one-dimensional case where trait evolution is restricted to only two directions, and would lead to greater complexity in evolutionary pathways and outcomes. However, the dimensionality of ℛ can, in some cases, be reduced by considering only the strain trait values that are of evolutionary interest, i.e., the set of points in ℛ that constitute a trade-off in *τ*-values for the considered host species (trade-off in compatibility). For example, if we only consider two host species and apply the Euclidean distance, then ℛ can safely be reduced to the one-dimensional case, see Supplementary material S6.

### (d) Conclusion and Outlook

The principal contribution of the work presented here is the ability to connect transmission parameters (the probability of infection, *τ*) to host species traits and predictors (important to viral sharing and pathogen-host combability), and the pathogen strain strategy. This makes the framework that was used here ideal for studying host-shift evolution of pathogens, for example for zoonotic virus. A challenge in applying our model to different scenarios, e.g., to specific host species and geographical areas, however, lies in the construction of the resource space (ℛ). One potential solution is to utilize data and method applied to the study of viral sharing (host-host) and pathogen-host association network data [13,25–27]. Specifically, one could use a parametric model, e.g., generalized additive mixed effect models (GAMMs), fitted to the network data, as a proxy for the *τ*- function and ℛ. This, together with other extensions, e.g., the addition of population dynamics, time-dependent contact rates, virulence, or within-host strain-competition, would extend the utility of our presented framework in the future.

## Data accessibility

Code for the model is available on GitHub at: https://github.com/PeterFransson/Evolution_and_Spillover_Project

## Authors’ contributions

P.F. and H.S. conceived the study and formulated the model. P.F. implemented and analyzed the model, and wrote the first draft of the manuscript. Both authors discussed the results and implications and revised the manuscript.

## Supplementary material

### S1 Stochastic simulations of the dual host-species system

To simulate realizations of our stochastic SIS model we employ the approximative hybrid method described in [31]. Here, we will briefly outline the most important aspects of the method, readers are referred to the original paper for a more detailed description. Let ***x***_*t*_ denote the current state of the system, i.e., the number of susceptible and the strain distribution (the collection of *I*_*i,z*_) for each species, at time *t*, and *α*_*e*_ is the propensity of an event *e* (Table S 1).

**Table S1.**
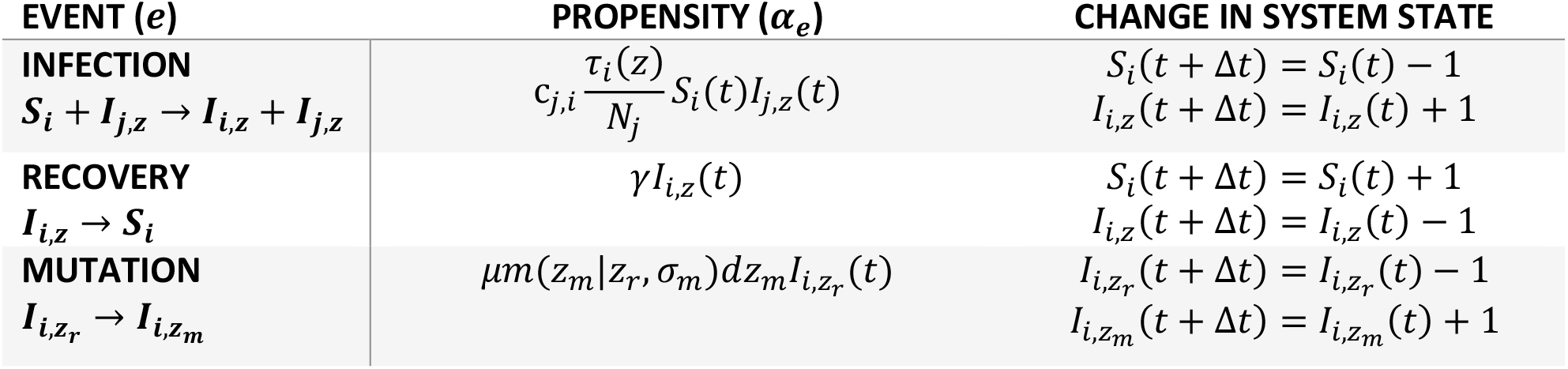
Equations for calculating the propensity for each event type and the resulting system state changes from time t to the time of the event at t + Δt. State variables and parameters are described in Table 1.

At each step of the method the next time step *h* and the events that occur during the time window [*t, t* + *h*] are determined. The next state, ***x***_*t*+*h*_, is determined according to the events that occurred (Table S 1). The method applies the exact SSA [30] for critical events (*e*_crit_) and the approximative tau-leaping method [32] for non-critical events (*e*_non−crit_). An event is critical if it impacts the size of a state variable, *I*_*i,z*_(*t*) at time *t*, which is less than a preset threshold *N*_*c*_, i.e., *I*_*i,z*_(*t*) ≤ *N*_*c*_; Otherwise, the event is said to be non-critical. Note, with this definition a mutation event is always considered a critical event. A time step for the non-critical events (tau-leaping), Δ*t*_non−crit_, is determined by finding the largest time step such that |*I*_*i,z*_(*t* + Δ*t*_non−crit_) − *I*_*i,z*_(*t*)| ≤ *εI*_*i,z*_(*t*) for all *I*_*i,z*_(*t*) > *N*_*c*_ [46]. Here *ε* (0.01 ∼ 0.1 [31]) is a preset error threshold. The time step for the critical events, Δ*t*_crit_, is determined by a draw from an exponential distributed random number with rate parameter 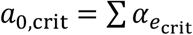. If Δ*t*_crit_ < Δ*t*_non−crit_ then a critical event is simulated according to the standard SSA. The final time step *h* = min (Δ*t*_crit_, Δ*t*_non−crit_) and the number of times a specific non-critical event occurred during 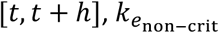, is determined by a draw from a Poisson distributed random number with mean 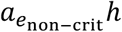. The system and time evolve until a preset time horizon *t*_end_ is reached, i.e., *t* ≥ *t*_end_. The final output is the evolution of ***x***_*t*_.

For our simulations we used *N*_*c*_ = 10 and *ε* = 0.1. We conducted simulations for the PIP outcomes: generalist (Figure S 1), evolutionary branching (Figure S 2), and bistable (Figure S 3). In all three cases we applied the following parameters: *N*_*a*_ = *N*_*b*_ = 10^3^, *x*_*a*_ = 0.2, *c*_*a,a*_ = *c*_*b,b*_ = 0.2, *c*_*a,b*_ = 0.12, *γ* = 0.1, *τ*_max_ = 1.0, *σ* = 0.05, *μ* = 0.01, *σ*_*m*_ = 0.003, *t*_end_ = 4500, and *x*_*b*_ = 0.27, 0.3, and 0.32 for the generalist, evolutionary branching, and bistable outcome, respectively. The initial condition (at *t* = 0) for all three cases was 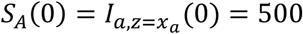 and *S*_*b*_(0) = *N*_*b*_. Thus, no individual in *b* was infectious and *z* = *x*_*a*_ was the only strain in circulation in the system at *t* = 0. This simulates a system when a reservoir host species population *a* comes into contact with a novel, fully susceptible, host species population *b*.

**Figure S1.**
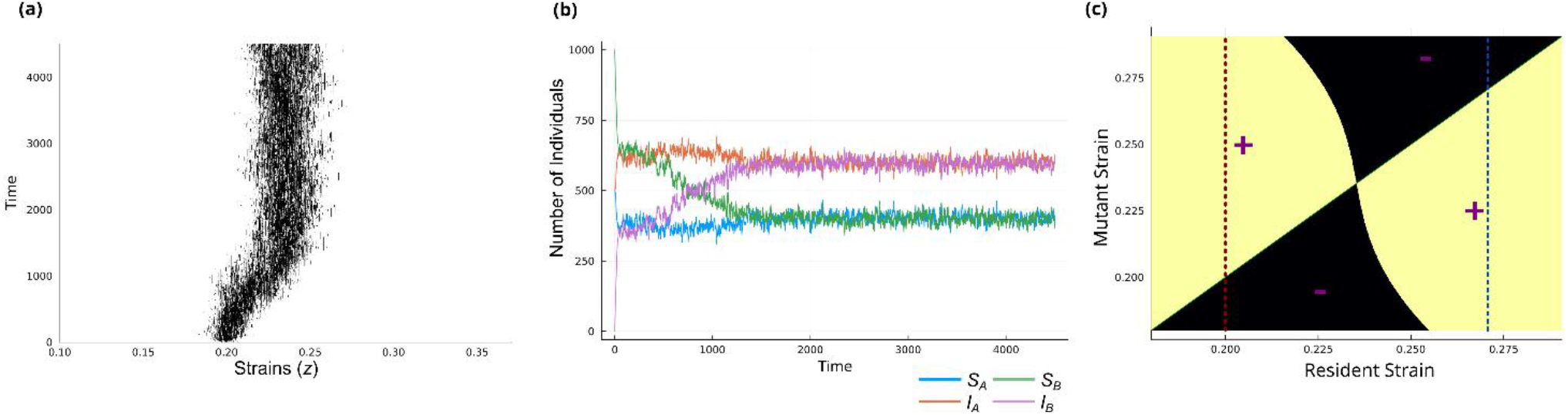
A trajectory generated from the stochastics SIS model for a generalist PIP outcome. The strain evolution is depicted in (a), where the black color at a particular xy-coordinate indicate the existence of at least one infectious individual at a specific time (y-coordinate), where the infection is caused by a specific strain (x-coordinate). More specifically the x-coordinate indicates the niche position of the strain. The evolution of the number of susceptible (S) and infectious (I) individuals in both host species population (host species a and b) are depicted in (b). Note, I_a_ and I_a_ are the sum of all infected individuals, i.e., I_a_ = ∑I_a,z_ and I_b_ = ∑I_b,z_. The corresponding PIP for the system is depicted in (c).

**Figure S2.**
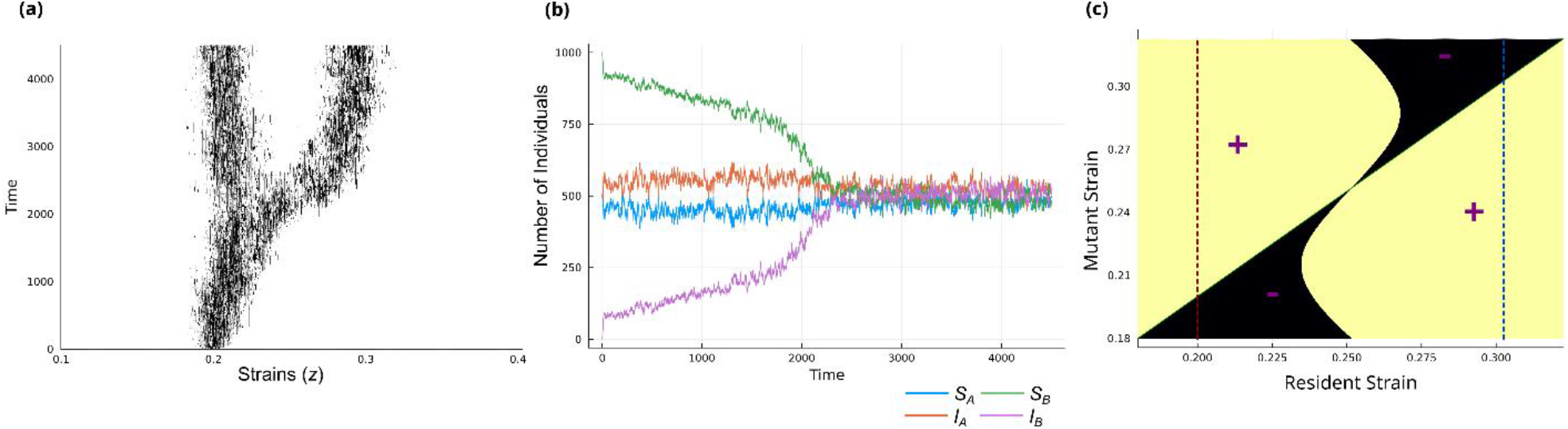
A trajectory generated from the stochastics SIS model for an evolutionary branching PIP outcome. The strain evolution is depicted in (a), where the black color at a particular xy-coordinate indicate the existence of at least one infectious individual at a specific time (y-coordinate), where the infection is caused by a specific strain (x-coordinate). More specifically the x-coordinate indicates the niche position of the strain. The evolution of the number of susceptible (S) and infectious (I) individuals in both host species population (host species a and b) are depicted in (b). Note I_a_ and I_a_ are the sum of all infected individuals, i.e., I_a_ = ∑I_a,z_ and I_b_ = ∑I_b,z_. The corresponding PIP for the system is depicted in (c).

**Figure S3.**
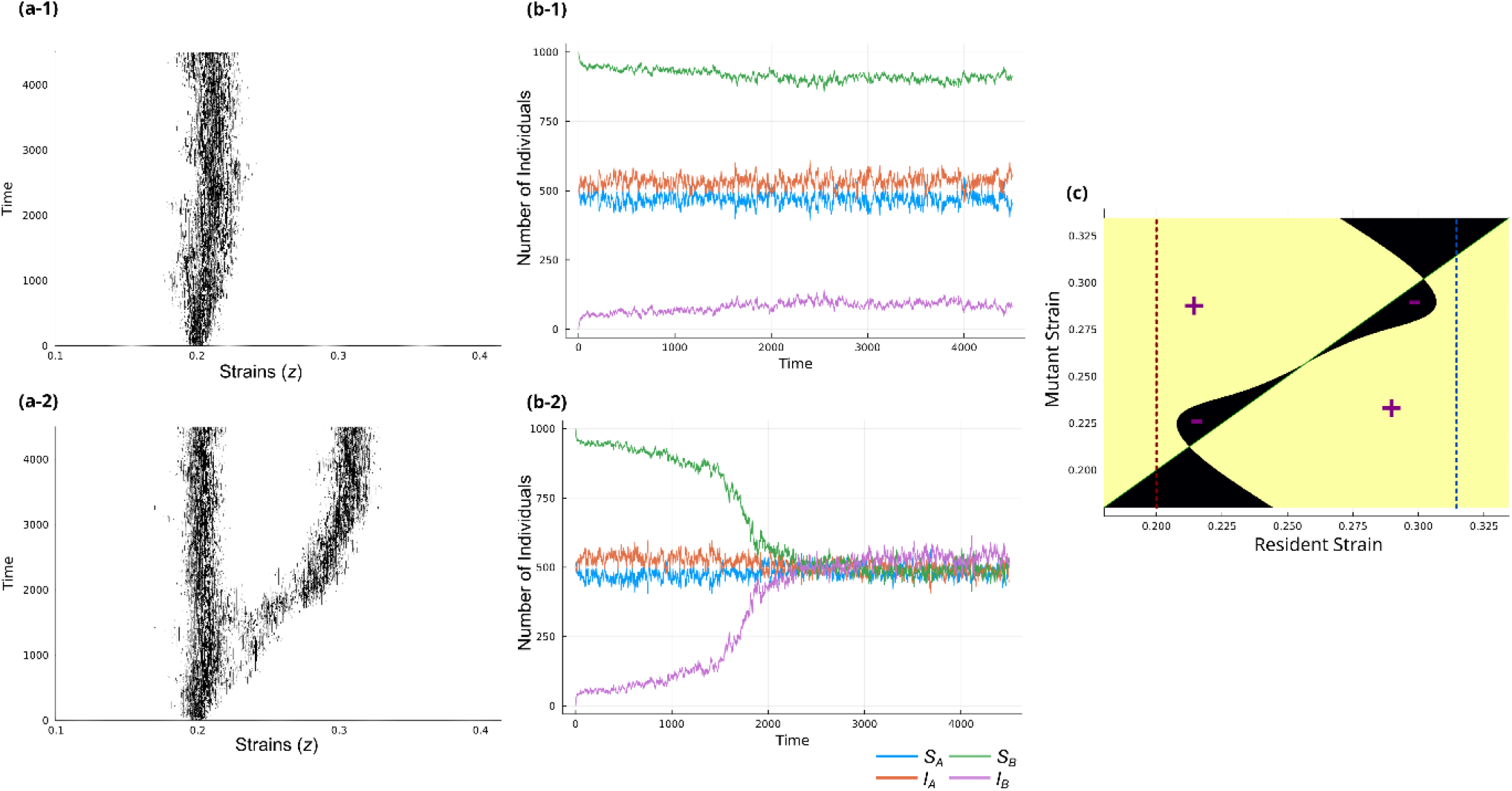
Two Trajectories generated from the stochastics SIS model for a bistable PIP outcome. The strain evolution is depicted in the a-panels (a-1 and a-2), where the black color at a particular xy-coordinate indicate the existence of at least one infectious individual at a specific time (y-coordinate), where the infection is caused by a specific strain (x-coordinate). More specifically the x-coordinate indicates the niche position of the strain. The evolution of the number of susceptible (S) and infectious (I) individuals in both host species population (host species a and b) are depicted in the B-panels (b-1 and b-2 I_a_ and I_a_ are the sum of all infected individuals, i.e., I_a_ = ∑I_a,z_ and I_b_ = ∑I_b,z_. The corresponding PIP for the system is depicted in (c). In the first trajectory (a-1 and b-2) the strain evolution gets “stuck” in the left most ESS (c) while in the second trajectory (a-2 and b-2) we observe an evolutionary branching event

### S2 Evolutionary invasion analysis

#### 2.1 Single-strain-multi-host model

For the evolutionary invasion analysis, we calculate the initial grows rate of a mutant strain with trait value *m* in the endemic state, determined by a resident strain with trait value *r*. To this end we need to express the dynamics of *r* to calculate the endemic state. The ODE that describes these dynamics is derived from a stochastic single-strain-multi-host version of our stochastic SIS model. Specifically, a version of the stochastic SIS model where we only consider infections being caused by one strain with trait value *r*, and mutation events are absent. The deterministic single-strain-multi-host ODE is then derived by mean-field approximation applied to the master’s equation of the stochastic model [35]. Let 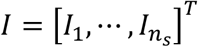 and 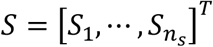 denote the vectors containing the number of infected and susceptible individuals within each of the *n*_*s*_ ∈ ℕ host species populations, respectively. The single-strain-multi-host ODE is then given by,

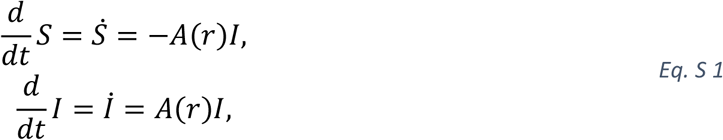

where *A*(*r*) is the system matrix,

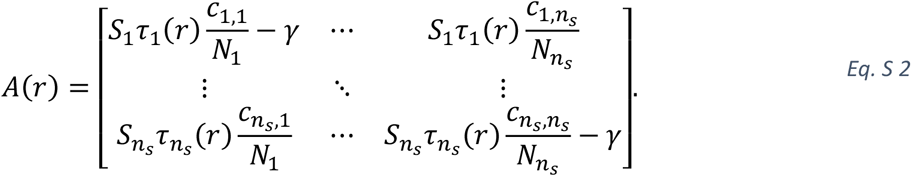

#### 2.2 Invasion fitness

From the single-strain-multi-host model (Eq. S 1), we derive the ODE which approximates the dynamics of an initial rare mutant strain with trait value *m*, in an environment set by the resident strain *r*. The system matrix for *m* has the same form as Eq. S 2, however it is depended on both *m* and *r*,

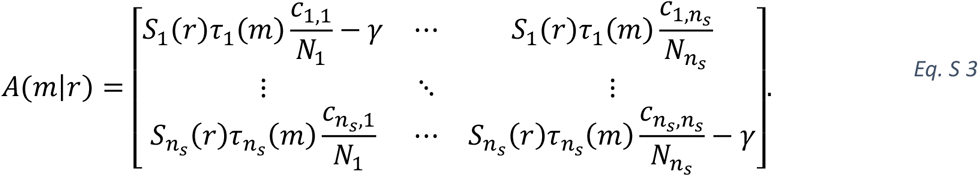

Here 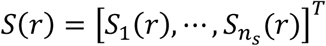 is the number of susceptible individuals for each species at the endemic state of *r*. The underlying assumption is that when *m* is rare then *S*(*r*) is approximately independent of *m* and does not change. Applying stability theory for linear systems of differential equations, we can determine if small perturbations, i.e., the introduction of *m* infected individuals, lead to an increase in *m* infected individuals –or if *m* goes extinct– by studying the eigenvalues of *A*(*m*|*r*). Specifically, *I*_*m*_ will go extinct if all eigenvalues of *A*(*m*|*r*) has a real part smaller than zero [47]. Thus, we define the invasion fitness as 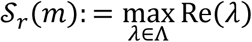, where Λ is the spectrum of *A*(*m*|*r*).

#### 2.3 Selection gradients

The selection gradient is given by the expression,

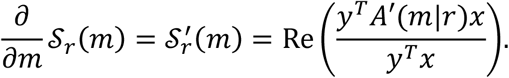

Here, *A*′(*m*|*r*) is the gradient matrix, *y* and *x* are the left and right eigenvector corresponding to 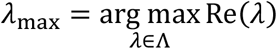, respectively, [48]. Note, the expression holds if the algebraic multiplicity of *λ*_max_ is one. The entries of the system matrix are continuous functions of the mutant trait value *m*. We get the gradient matrix by differentiating *A*(*m*|*r*) with respect to *m*,

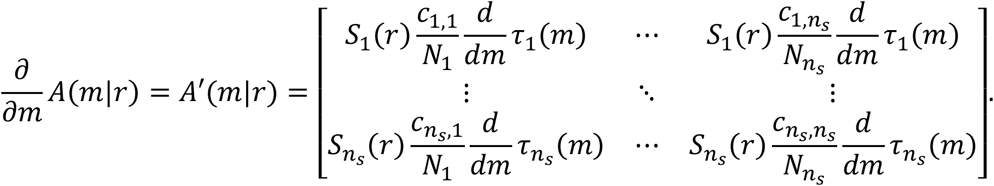

### S3 The dual host-species system

#### 3.1 Evolutionary invasion analysis

In the special case when we only consider two host species, then the system matrix, *A*(*m*|*r*) (Eq. S 3) is a 2 × 2 matrix. In this case *S*_*r*_(*m*) is given by the following expression,

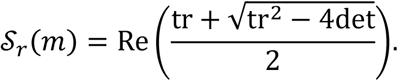

Here tr and det are the trace and determinant of *A*(*m*|*r*) respectively. We get an expression for the selection gradient by differentiation of *S*_*r*_(*m*),

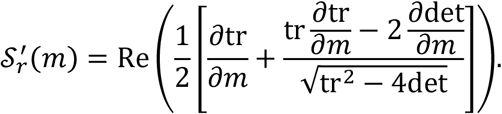

The second order derivative of *S*_*r*_(*m*) is given by,

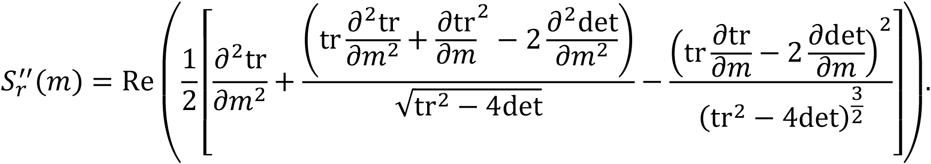

### 3.2 Parameters to determine the pairwise invasibility plot (PIP)

In this section we will reparametrize the dual host-species system and determine the parameters necessary to draw PIPs for our system. Recall that for any pair *r, m* ∈ ℛ we need to 1) find the endemic state of the resident strain *r* and 2) calculate the sign of the invasion fitness, *S*_*r*_(*m*), of the mutant strain *m*. Thus, we will determine the parameters needed to calculate these two.

First, we make a variable substitution, specifically, we introduce *i*_*j*_ = *I*_*j*_/*N*_*j*_ ∈ [0,1] and *s*_*j*_ = *S*_*j*_/*N*_*j*_ = 1 − *i*_*j*_ ∈ [0,1], the proportion of infected and susceptible individuals in population *j* ∈ {1,2}, respectively. Note, *di*_*j*_/*dt* = 1/*N*_*j*_ × *dI*_*j*_/*dt*, or in matrix form,

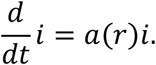

Where, *i* = [*i*_1_, *i*_2_]^*T*^ and,

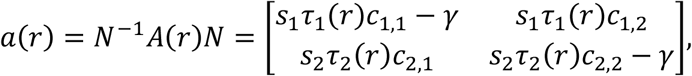

where,

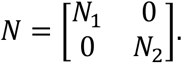

Similarly to *a*(*r*), we can define *a*(*m*|*r*) = *N*^−1^*A*(*m*|*r*)*N*. Since the matrices *A*(*m*|*r*) and *a*(*m*|*r*) are similar, they share the same eigenvalues and thus *a*(*m*|*r*) has the same stability as *A*(*m*|*r*). We find the endemic state *s*(*r*) = [*s*_1_(*r*), *s*_2_(*r*)]^*T*^, necessary for determining *a*(*m*|*r*), by solving the nonlinear system *a*(*r*)*i* = 0. Thus, the PIP is independent of *N*_1_ and *N*_2_ for the system. Note, this variable substitution works for the multi-host case.

Let 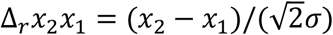, where *x*_1_ and *x*_2_ are the positions of the two host species in the resource space. We can then define the distance between points in resource space in terms of Δ_*r*_*x*_2_*x*_1_, e.g., Δ_*r*_*x*_1_*z* = −Δ_*r*_*zx*_1_ = −*δ*_*z*_Δ_*r*_*x*_2_*x*_1_ and Δ_*r*_*x*_2_*z* = (1 − *δ*_*z*_)Δ_*r*_*x*_2_*x*_1_ are the distances between a given strain *z*, and *x*_1_ and *x*_2_, respectively. Here *δ*_*z*_ ≔ (*z* − *x*_1_)/(*x*_2_ − *x*_1_). Note, the only strains of interest are *δ*_*z*_ ∈ [0,1]. With these notations we can rewrite the ODE for the single-strain-two-species system,

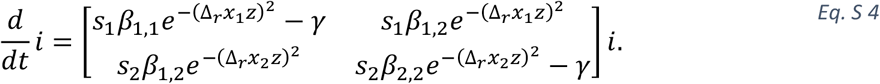

Here, *β*_*i,j*_ = *τ*_max_*c*_*i,j*_. We can further simply the system by factorizing *γ* and introducing the parameter *R*_0,*i,j*_ = *β*_*i,j*_/*γ* (basic reproduction ratio),

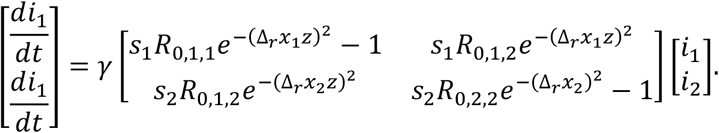

We find the endemic state by solving the system of nonlinear equations,

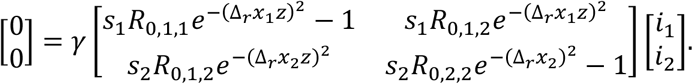

Multiplying by *γ*^−1^ on the r.h.s. and l.h.s., we get the equivalent equation,

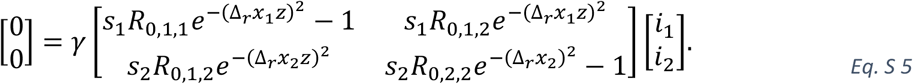

Thus, the necessary parameters to determine the endemic state of the two-host-species system are:

Remember that the dynamics of a rare mutant strain *m* is determined by the system matrix *a*(*m, r*). Using the previously introduced notations and factorizing *γ, a*(*m, r*) can be rewritten,

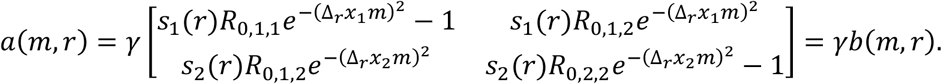

We calculate *S*_*r*_(*m*) from the eigenvalues of *a*(*m, r*). We get the eigenvalues by solving,

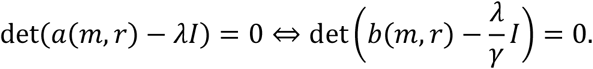

This implies that, if *λ*_*B*_ is an eigenvalue of *b*(*m, r*), then *λ*_*A*_ = *γλ*_*B*_ is an eigenvalue of *a*(*m, r*). Since *γ* > 0, then sign(*λ*_*A*_) = sign(*λ*_*B*_), i.e., the stability of *a*(*m, r*) can be determined by studying *b*(*m, r*). Consequently, no additional parameters are needed to calculate the sign of *S*_*r*_(*m*) and the PIPs for the two-host species system are determined by the four parameters:

*R*_0,1,1_, *R*_0,2,2_, *R*_0,1,2_, Δ_*r*_*x*_2_*x*_1_. Out of convenience, we substitute *R*_0,1,2_ for the parameters *C*_ratio_ in our simulations. We define *C*_ratio_ as,

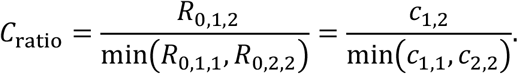

#### 3.3 Expressions for the endemic state

In this section we will derive expressions for the endemic state, *S*(*r*) = [*S*_1_, *S*_2_]^*T*^, for the dual host-species-one-strain model. The endemic state is obtained by finding solutions to the equation *B*(*r*)*I* = 0. For any non-trivial solution to the homogeneous equation, the matrix,

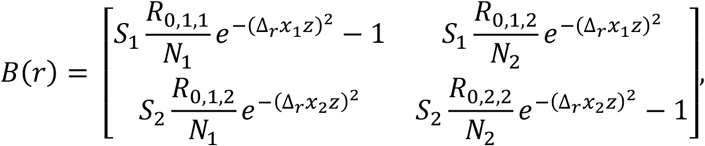

needs to be singular, i.e.,

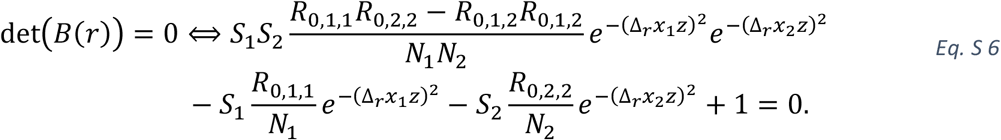

Additionally, the solutions (*X*^∗^) to the homogeneous equation, i.e., *B*(*r*)*X*^∗^ = 0, can be expressed as,

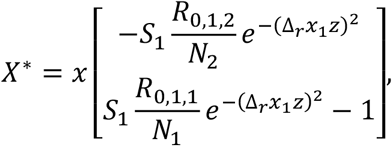

where *x* ∈ ℝ. We have two additional conditions: i) *X*^∗^ = [*I*_1_ *I*_2_]^*T*^ and ii) *I*_*i*_ = *N*_*i*_ − *S*_*i*_ ∀*i* ∈ {1,2}. This implies,

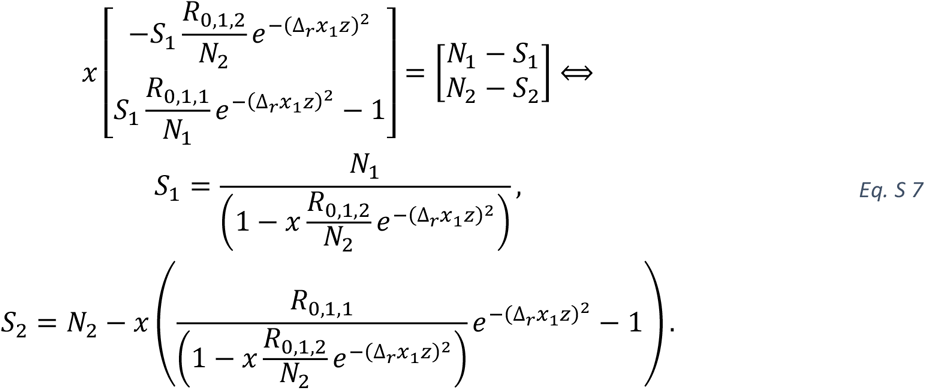

Thus, we have expressions for *S*_1_ and *S*_2_ in terms of *x*. We can plug these into Eq. S 6, to obtain a final equation expressed in terms of *x*, see below.

Before we derive the expressions for the endemic state, we will derive constrains on the variable *x*. Note, we are only interested in the solutions obeying the condition 0 < *S*_*i*_ < *N*_*i*_, ∀*i* ∈ {1,2}. For the condition 0 < *S*_1_ < *N*_1_ we get,

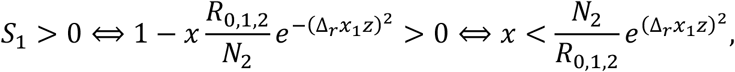

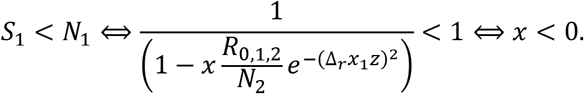

Thus, *x* < 0. For the second condition, 0 < *S*_2_ < *N*_2_, we have

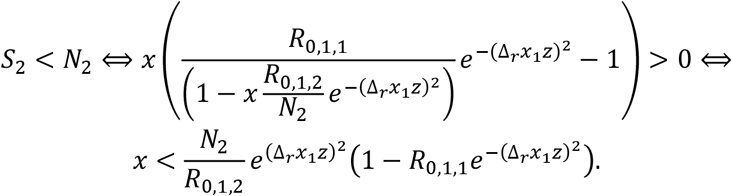

Additionally,

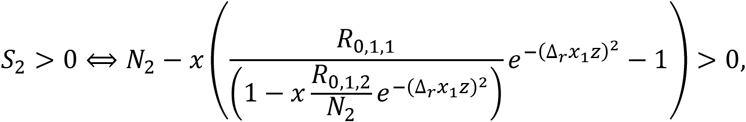

Which is equivalent to,

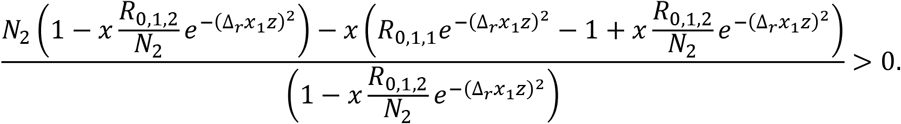

Since the denominator is always positive (a consequence from the condition *x* < 0), the numerator needs to be positive for the inequality to be satisfied,

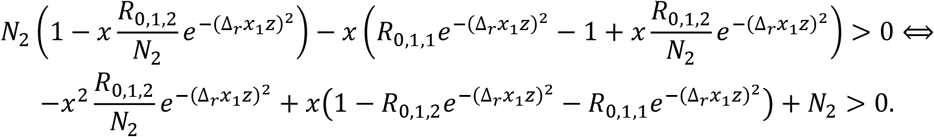

The expression on the left-hand side is a quadratic polynomial. Since *N*_2_ > 0 and the coefficient in front of *x*^2^ is negative, the corresponding quadratic equation will have a positive and a negative real root. Thus, the left-hand side is > 0 when *x* is between the two roots. We are only interested in the case when *x* < 0, thus,

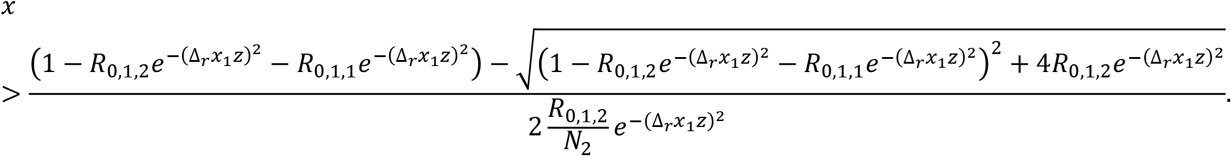

Thus, the final interval of feasible solutions is,

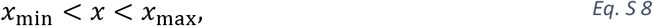

where,

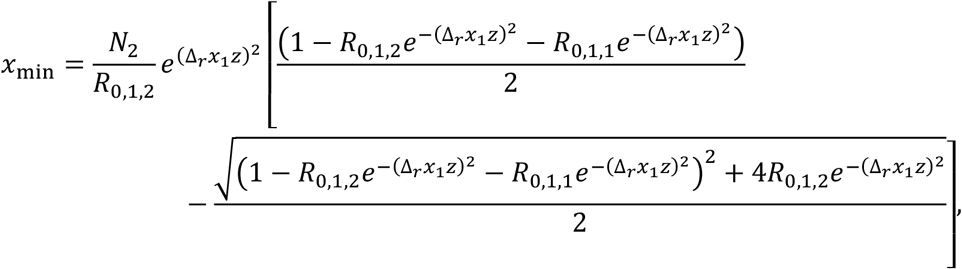

and

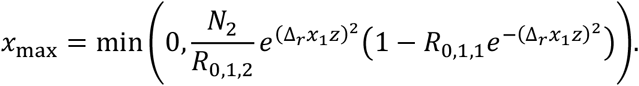

We will now derive the expression for the endemic state of the two-host-species-one-strain system. We can simplify Eq. S 6 by introducing new parameters (*a*_eq_, *b*_eq_, *c*_eq_),

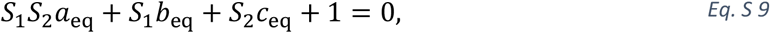

where,

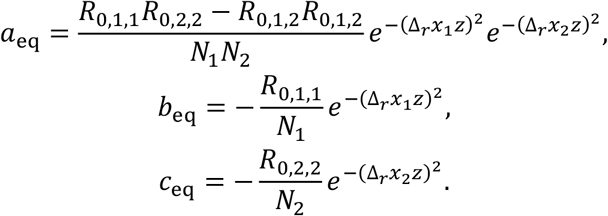

Now we substitute with the expressions for *S*_1_ and *S*_2_ (Eq. S 7) in Eq. S 9 to derive an expression in terms of *x*. Observe that the product *S*_1_*S*_2_ yields a fraction with a denominator equal to

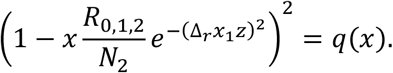

More specifically,

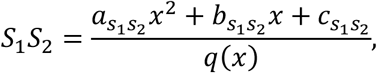

where,

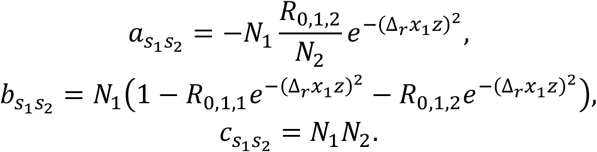

Note, *q*(*x*) > 0 due to the imposed condition *x* < 0. We want to express each term in Eq. S 9 as a fraction with *q*(*x*) as the denominator. To this end we have to express *S*_1_, *S*_2_, and 1 as fractions:

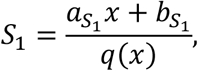

where,

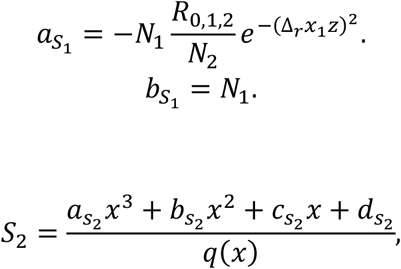

where,

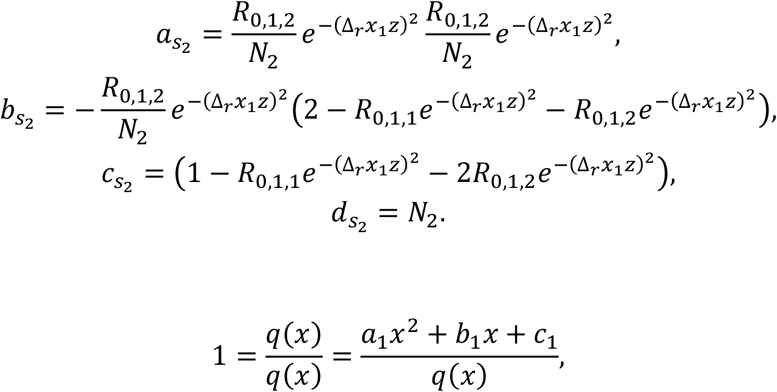

where,

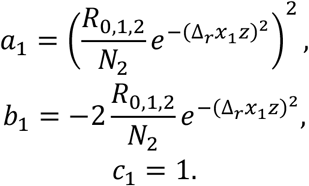

Since all terms share the same denominator, we sum the numerators from the terms to obtain a cubic polynomial,

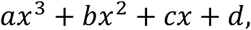

Where,

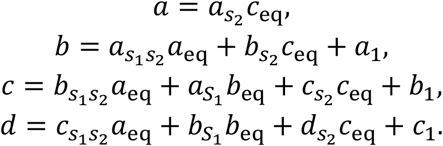

Note, *S*_1_*S*_2_*a*_eq_ + *S*_1_*b*_eq_ − *S*_2_*c*_eq_ + 1 = 0 ⟺ *ax*^3^ + *bx*^2^ + *cx* + *d* = 0. To obtain the real solutions of the cubic equation, *x*^∗^, we rewrite the equation to its depressed cubic form,

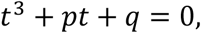

where,

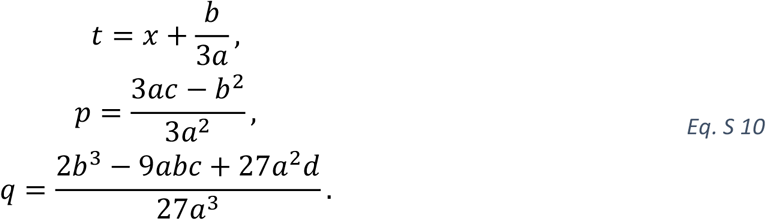

The number of real solutions, and their expression, depends on the sign of the expression *q*^2^/4 + *p*^3^/27 [49]. If *q*^2^/4 + *p*^3^/27 = 0 and *p* = 0, then the depressed cubic form has a triple root 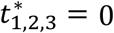 . If *q*^2^/4 + *p*^3^/27 = 0 and *p* ≠ 0, then then depressed cubic has three real solutions, one double root 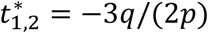 and 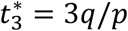. If *q*^2^/4 + *p*^3^/27 > 0 then depressed cubic has one real root (Cardon’s formula),

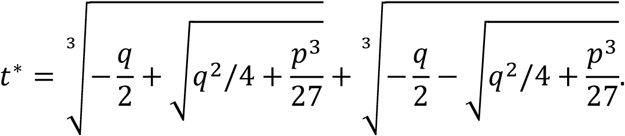

If *q*^2^/4 + *p*^3^/27 < 0, then the depressed cubic has three, distinct, real solutions,

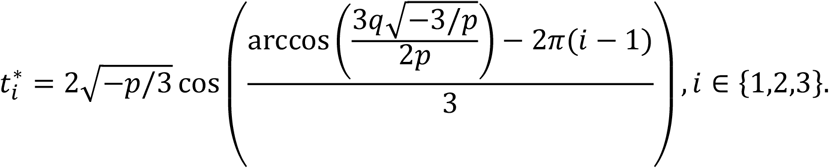

We obtain the final endemic state *S*(*r*) by using Eq. S 10 to obtain *x*^∗^, checking condition Eq. S 8 to identify the feasible solutions, and applying Eq. S 7.

### S4 Trait evolution plot

**Figure S4.**
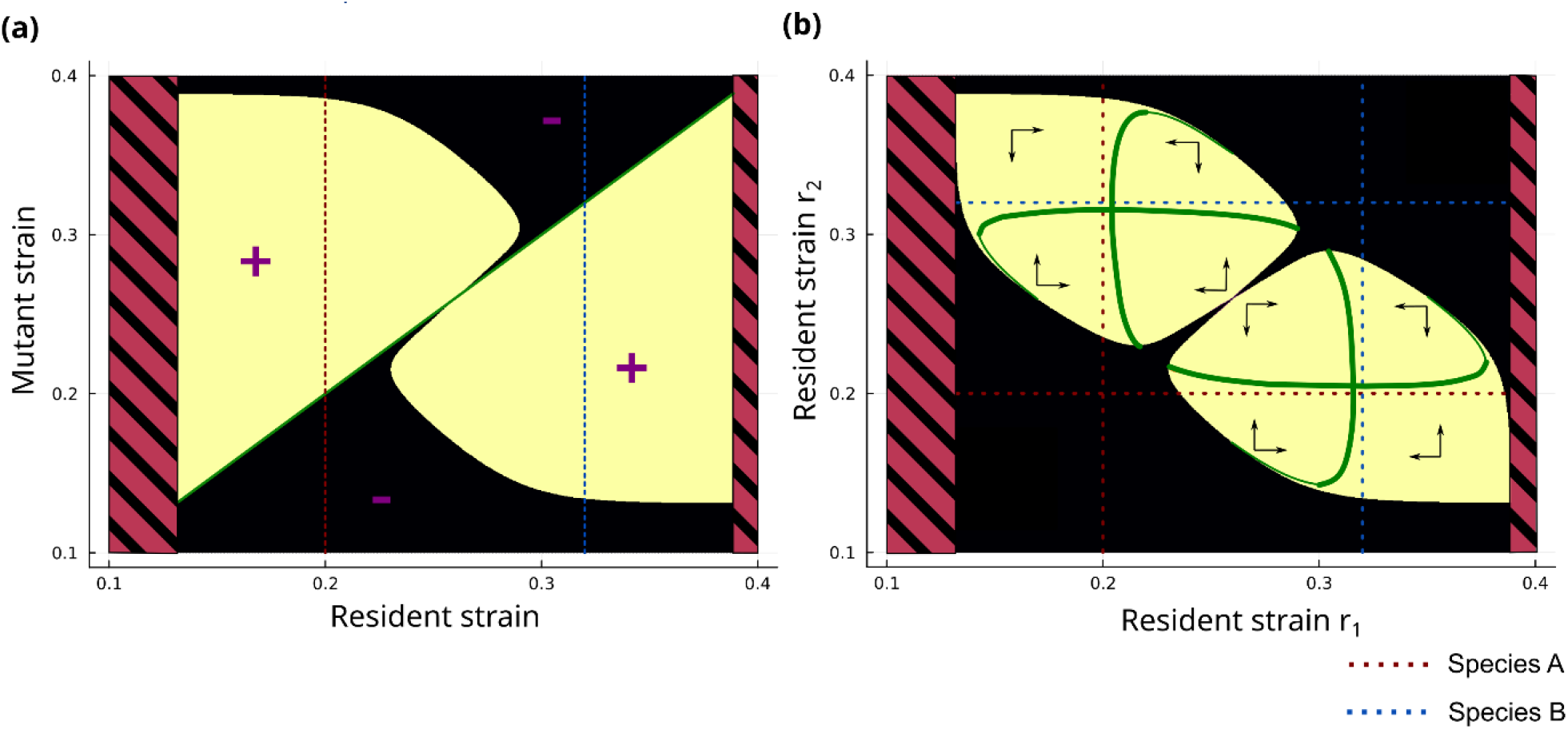
Evolutionary branching system. Similarity between the two-host species (A and B) is graphically depicted as the distance between the red and blue dashed lines in (a) and (b). In the evolutionary branching outcome, the pairwise invasibility plot (a) has an evolutionary branching point. The initial monomorphic strain population evolves toward this point at which the population undergo disruptive evolution and two coexisting strain trait values, r_1_ and r_2_, emerges, i.e., we get a coexisting dimorphic strain population. The trait evolution plot (b) shows the region of possible coexisting resident strain trait values (light shaded area). The direction of evolutionary change is indicated by arrows (invasion cones). Green lines are isoclines where the selection gradient vanishes. Thick and thin lines correspond to a fitness maximum and minimum, respectively. The intersection between the isoclines shows an evolutionary stable strategy for the dimorphic population. The figures were generated using the following parameter values: R_0,a_ = R_0,b_ = 2.55, c_ratio_ = 0.71, and Δ_r_x_a_x_b_ = 1.75.

### S5 Additional interspecific contact intensity figures

**Figure S5.**
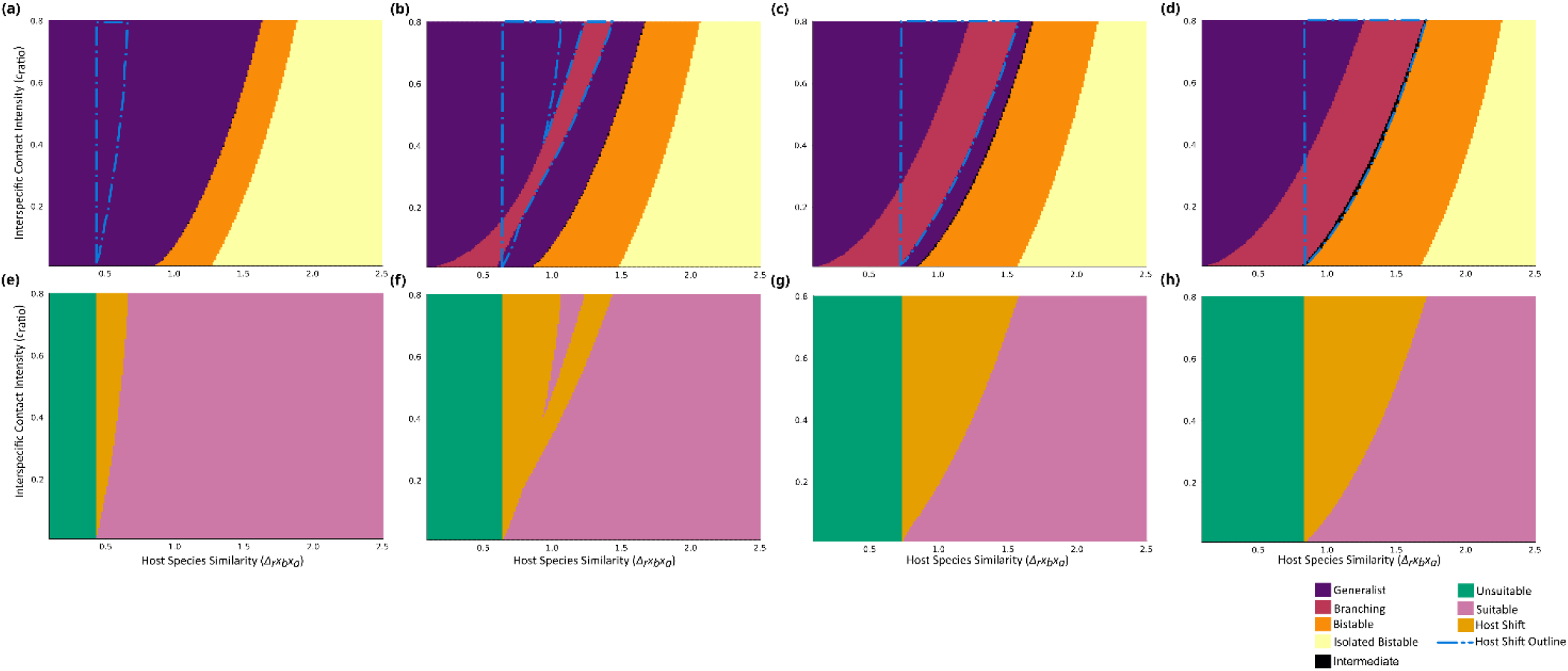
The impact of R_0,a_ and R_0,b_ on evolutionary outcome and evolutionary host shifts (EHS) events. The color of the top heatmaps (a-d) indicates the evolutionary outcome (Generalist, (evolutionary) Branching, Bistable, Isolated Bistable, and Intermediate outcomes) for different combinations of host species similarity (Δ_r_x_b_x_a_) and Interspecific contact intensity (c_ratio_). The color of the bottom heatmaps (e-h) indicates if a Δ_r_x_b_x_a_ and c_ratio_ combination describes a system which is: not an evolutionary host shift scenario (EHSS) (Unsuitable), i.e., a parameter combination not fulfilling necessary conditions for a EHS to occur. EHSS, but EHS did not occur (Suitable). EHHS and EHS occurred (Host Shift). Note the dashed curve in (a)-(d) indicates the corresponding outline of the Host Shift region in (e)-(h). The heatmaps where generated using R_0,b_ = 1.2 (a, e), R_0,b_ = 1.5 (b, f), R_0,b_ = 1.7 (c, g), and R_0,b_ = 2.5 (d, h). For all heatmaps R_0,a_ = 2.0.

### S6 Extension to Higher-Dimensional Resource Spaces

We have thus far restricted the dimension of the resource space to one. For the extension to higher dimensions, we introduce the following definitions: let ℛ = ℝ^*d*^ denote the *d*-dimensional resource space, *d*(*x, y*) is the Euclidean distance between the points *x, y* ∈ ℛ.

For a system with two host species, species *a* and *b*, we can always transform the *d*-dimensional recourse space into the one-dimensional case. Let *x*_*a*_, *x*_*b*_ ∈ ℛ denote the representation of *a* and *b*, respectively. As previously mentioned, *τ*(*x, z*) represents the compatibility between a host species *x* ∈ ℛ and a stain *z* ∈ ℛ. From a multi-objective optimization perspective, the *τ*-values 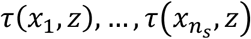 for a finite set of host species 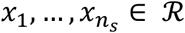 can be viewed as a set of objective functions, from the pathogen’s perspective. We are only interested in the stain strategies which results in a trade-off in the *τ*-values, i.e., strains that cannot increase their *τ*-value in one host species without losing compatibility (i.e., lower *τ*-value) in another. From the multi-objective optimization perspective, these strain strategies are referred to as Pareto-optimal solutions [50]. The set of all Pareto-optimal solutions are called the Pareto front and this set represents the collection of strain strategies which are of evolutionary interest. Since *τ*(*x, z*) is a monotonically decreasing function of *d*(*x, z*), we maximize *τ*(*x, z*) by minimizing the distance between *x* and *z*. Let *L* = {*x* ∈ ℛ: *x* = *k*(*x*_*b*_ − *x*_*a*_) + *x*_*a*_, *k* ∈ ℝ} denote the line that passes through *x*_*a*_ and *x*_*b*_, and 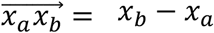. Let 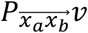 denote the orthogonal projection of a vector *v* on to 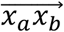. Let *z* denote an arbitrary strain. We can decompose the vector 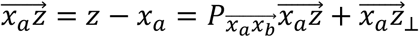, where 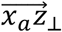 is a vector, orthogonal to 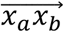. From the law of cosines, we get 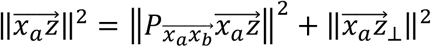 Thus 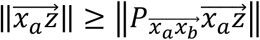, where we have an equality only if *z* ∈ *L*. This implies that the orthogonal projection of the point *z* on to 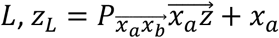, is better adapted to *a*, i.e., *τ*(*x*_*a*_, *z*_*L*_) > *τ*(*x*_*a*_, *z*). Since this also holds true for *x*_*b*_, then *z*_*L*_ is better adapted to both *a* and *b*. It directly follows that for every strain, *z*^∗^, not contained in *L*, there exist at least one strain *z* in *L* which is better adapted to both *a* and *b*. Mathematically expressed, 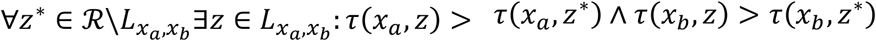 . Thus, *L* –more accurately the line segment between *x*_*a*_ and *x*_*b*_– represents the Pareto front and we can make the following variable substitution *z* = *z*^∗^*x*_*b*_ + (1 − *z*^∗^)*x*_*a*_, to transform the *d*-dimensional recourse space into the 1-dimensional case. Here, *z*^∗^ is the new strain strategy and ℛ^∗^ = ℝ is the new resource space. Similar arguments can be made for systems with more than two host species.

A limitation to this extension is that we assume that all phenotypes and covariates, represented by the different axis in ℝ^*d*^, have comparable units (i.e., one-unit change in one axis is equivalent to one-unit change in another) and are uncorrelated. However, the underlying covariates can be scaled or normalized to address the issue of noncomparable units. If the set of covariates are linearly depended then we can find a linear map that changes the basis of the space to a linearly independent basis, e.g., by applying principal component analysis (PCA). If nonlinear relationships exist between the covariates, or if the covariates are noncontinuous (e.g., categorical covariates), other methods could be applied, e.g., nonlinear principal components analysis [51].

